# Polyploidy-associated paramutation in Arabidopsis is determined by small RNAs, temperature, and allele structure

**DOI:** 10.1101/2020.10.21.348839

**Authors:** Heinrich Bente, Andrea M. Foerster, Nicole Lettner, Ortrun Mittelsten Scheid

## Abstract

Paramutation is a form of non-Mendelian inheritance in which the expression of a paramutable allele changes when it encounters a paramutagenic allele. This change in expression of the paramutable alleles is stably inherited even after segregation of both alleles. While the discovery of paramutation and studies of its underlying mechanism were made with alleles that change plant pigmentation, paramutation-like phenomena are known to modulate the expression of other traits and in other eukaryotes, and many cases have probably gone undetected. It is likely that epigenetic mechanisms are responsible for the phenomenon, as paramutation forms epialleles, genes with identical sequences but different expression states. This could account for the intergenerational inheritance of the paramutated allele, providing profound evidence that triggered epigenetic changes can be maintained over generations. Here, we use a case of paramutation that affects a transgenic selection reporter gene in tetraploid *Arabidopsis thaliana*. Our data suggest that different types of small RNA are derived from paramutable and paramutagenic epialleles. In addition, deletion of a repeat within the epiallele changes its paramutability. Further, the temperature during the growth of the epiallelic hybrids determines the degree and timing of the allelic interaction. The data further make it plausible why paramutation in this system becomes evident only in the segregating F2 population of tetraploid plants containing both epialleles. In summary, the results support a model for polyploidy-associated paramutation, with similarities as well as distinctions from other cases of paramutation.

**AUTHOR SUMMARY:** In 1866, Gregor Mendel formulated the general principles of inheritance based on crossing experiments with pea plants. Curiously, in 1915, the progeny from crossing pea plants with a regular and a “rogue” leaf phenotype was lacking the expected segregation and recovery of the regular phenotype. This discovery was one of the first observations of non-Mendelian genetics and later demonstrated for more traits in other plants and termed paramutation. Paramutation is due to the epigenetic switch of an active gene to a silenced version which is then maintained in the inactive state in later generations. This demonstrates that acquired epigenetic changes can become permanent. Despite its early observation and numerous studies, mainly in maize and tomato, it is barely understood how paramutation is established and which parameters influence the process. We investigated a case of paramutation in *Arabidopsis thaliana*, crossing plants with genetically identical but epigenetically different alleles that result in resistance or sensitivity to an antibiotic in the growth medium. Paramutation did not become manifest immediately but only in the progeny of the hybrids, and only in plants with a doubled chromosome set. These features make this paramutation distinct from other cases. Our studies revealed several parameters that influence paramutation: an important role for sRNAs to initiate silencing, the sequence of the allele itself, the environmental conditions during growth of the hybrids, the developmental stage, and the copy number ratio between the alleles.

## INTRODUCTION

Paramutation is the term for a specific interaction between alleles of a gene, in which one paramutable allele becomes heritably modified by encounter with a paramutagenic allele. It violates the Mendelian rule of independent segregation, as the paramutated allele is maintained in the new state and even can acquire paramutagenicity itself. This led to its description as para-(resembling)-mutation or somatic gene conversion by the pioneers working on maize and tomato in the 1950’s [1–3]. Even earlier genetic experiments with peas [4] are now interpreted as evidence of paramutation [5]. Numerous cases of paramutation have been reported in several species including metazoans and plants [reviewed in 6, 7, 8], indicating that paramutation is not just a bizarre exception but may be a general phenomenon among eukaryotes [9].

Paramutation has an epigenetic basis and does not change the DNA sequence of participating alleles. Instead, specific, newly acquired epigenetic states can be installed and maintained over generations independent of the trigger of the change. It is likely the best example for true transgenerational inheritance of defined epigenetic changes connected with a specific trigger, namely the encounter with the paramutagenic allele. Although many principles of epigenetic regulation are now known, the molecular basis of paramutation is still not well understood. Genetic screens, mainly in maize, have identified several genes whose function is necessary for establishment and/or maintenance of paramutation, e.g. *“required to maintain repression”* (*rmr*) [10] or *“mediator of paramutation”* (*mop*) [11]. The nature of their gene products [reviewed in 12] suggested that paramutation requires components of the RNA- directed DNA methylation (RdDM) pathway [reviewed in 13, 14, 15], which is responsible for silencing repetitive sequences via small RNAs (sRNAs) that guide the RNA-induced silencing complex (RISC) to corresponding genomic loci, leading to DNA methylation and the formation of heterochromatin. However, the majority of RdDM targets are not paramutable, suggesting that additional factors, such as *cis*-acting determinants [16] or other *trans*-acting elements are involved. To identify these, additional paramutation systems will be informative, especially those that differ in penetrance, frequency, and stability of the epigenetic change.

Paramutation has also been demonstrated in *Arabidopsis thaliana* [17]. It involves epialleles of a transgene that, in an active state, expresses hygromycin phosphotransferase (HPT) conferring resistance (R epiallele) to hygromycin. The inactive state confers sensitivity (S epiallele) to the antibiotic. *In vitro* culture of protoplasts from diploid plants with an R epiallele and subsequent regeneration resulted in several tetraploid plants, some of which were hygromycin-resistant while others had spontaneously acquired a silent epiallele. These silenced plants acquired DNA methylation and histone marks characteristic of silenced chromatin [18, 19], no longer transcribed the *HPT* gene [17], and rendered plants hygromycin-sensitive. Diploid R and S derivatives were generated by backcrossing to diploids, and progeny that were homozygous for either R or S alleles inherited their respective expression state and phenotype over numerous generations, in both, diploid and tetraploid lines. Molecular analysis confirmed that the sequence of the R and S allele were identical, and the *HPT* gene is inserted in an intergenic region far from transposons. Therefore, it was concluded that the R and S alleles differ in their epigenetic state and are true epialleles [20]. A mutant screen scoring for reactivation of the silent epiallele led to identification of the *trans*-acting mutants *ddm1* and *hog1*, confirming the importance of DNA and histone methylation [18]. It also demonstrated that *cis*-acting mutations with structural variants of the epiallele (RΔ) led to high levels of HPT transcripts and hygromycin resistance [20]. Together these data demonstrated that the S allele results from epigenetic silencing of the *HPT* gene.

Crossing diploid R and S plants resulted in 100% hygromycin-resistant F1 plants, and selfed F1 plants produced F2 progeny with close to 75% (3:1) resistant plants, as expected for a dominant trait and confirming independent segregation of the epialleles. However, while F1 hybrids from crosses between tetraploid R and S plants were 100% resistant, their F2 progeny contained significantly fewer resistant plants than the expected 97% (35:1), although there was variation between independent tetraploid F2 populations. These data argued against a genetic determinant of the lost resistance and suggested that epigenetic components were interfering with the independence of segregation, similar to paramutation [17]. However, there were interesting differences from the classical paramutation cases. While the loss of gene expression was visible in maize and tomato in the F1 hybrid [21] of these diploid species, the phenotype indicating paramutation in the R/S interaction was restricted to the F2 generation of tetraploid Arabidopsis hybrids only. These data led to the expectation that this system’s late onset of the interaction and its requirement for polyploidy could provide more insight into the mechanisms that determine the occurrence and degree of paramutation.

Here, we investigated the role of the epialleles’ structure, the connection with transcript abundance, and the potential for secondary paramutation. We examined the effect of temperature during growth of the F1 hybrids, and profiled small RNAs associated with the epialleles in seedlings and flower buds of diploid and tetraploid plants. We show that the degree of paramutation depends on environmental conditions, developmental stage, and an interplay between the structure of the epialleles and the nature of the small RNAs associated with them. Based on these data, we present a model that explains the role of copy number in the epiallelic interaction in polyploid plants.

## RESULTS

### Nature and nomenclature of the epialleles involved in paramutation

The phenotype revealing paramutation in Arabidopsis is based on expression or silencing of the hygromycin phosphotransferase (*HPT*) gene conferring resistance to the antibiotic hygromycin. A construct with *HPT* under control of the CaMV 35S promoter was used for PEG-mediated protoplast transformation [22] and inserted in an intergenic region of chromosome three in the Arabidopsis genome. Plasmid copies recombined before integration, resulting in an insert consisting of a vector fragment upstream of the promoter and the *HPT* open reading frame, a partially deleted CaMV 35S terminator followed by a second copy of the vector and the CaMV 35S promoter, and approximately 500 bp non-coding non-plant carrier DNA used during the protoplast transformation. This structure has been previously described [17, 20]; a schematic illustration of all variants used in this study (considering the transcription direction opposite to the reference genome) is shown in Fig 1A. Active and silenced epialleles have identical DNA sequence and are hereafter referred to as R (resistant) or S (sensitive). The “empty” insertion site in the wild type is termed W. Diploid and tetraploid genotypes will be called RR/SS/WW and RRRR/SSSS/WWWW, respectively. An overview of the origin and crosses of each epiallele and the respective mutants is illustrated in Fig 1B.

**Figure 1.**
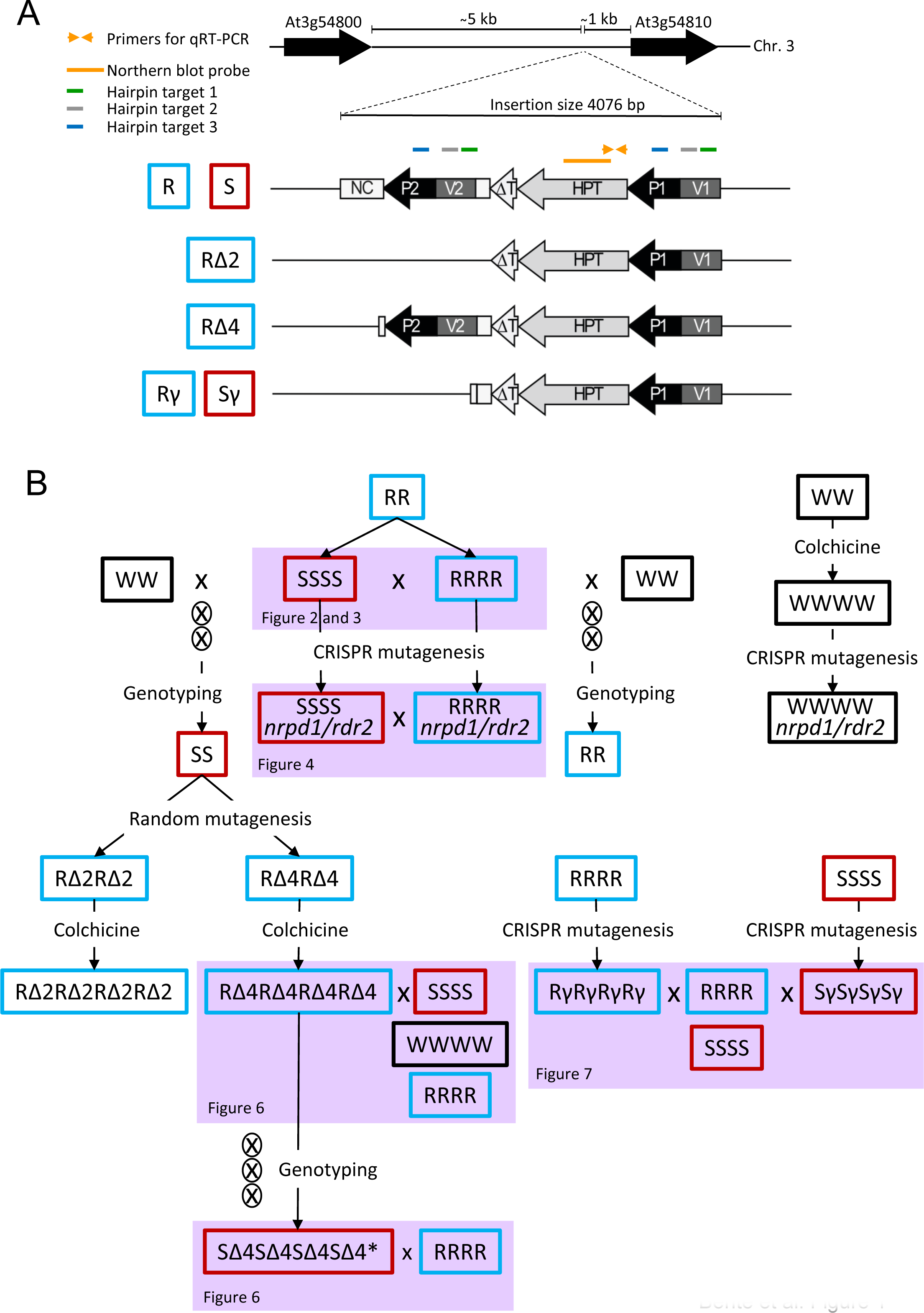
Overview over structure of epialleles and origin of plant material. (A) Location of the epialleles in an intergenic region on chromosome three (top), the different structural variants below, and the location of primers and probes. HPT: ORF of hygromycin phosphotransferase; ΔT: incomplete CaMV 35S terminator; P1, P2: CaMV 35S promoter; V1, V2: vector plasmid sequence; NC: non-coding sequence; S: silent epiallele rendering plants sensitive (red); R: active epiallele rendering plants resistant (blue) to hygromycin; Δ, γ: deletion derivatives (obtained by random or CRISPR mutagenesis, see text for details and B). (B) Pedigree of lines. S, R, Δ, γ as in (A), W: wild type allele without insert (black). All lines are derived from the same, initially hygromycin-resistant homozygous transgenic line RR. RRRR and SSSS are two independent lines obtained by spontaneous polyploidization during somatic cloning. 2 letters: diploids; 4 letters: tetraploids; x: crossing between different parent plants; 955 progeny by self-fertilization; colchicine: generation of tetraploids, *: converted by paramutation. Genotyping by PCR confirmed homozygosity of the alleles. Links to other figures refer to the use of the material in different context.

### Paramutation depends on temperature during growth of tetraploid F1 hybrids

Plants homozygous for either R or S inherit hygromycin resistance or sensitivity as stable traits. Hybrids obtained by crossing diploid plants and selfing them resulted in F2 progeny with the expected 3:1 ratio, regardless whether R was combined with S or with the wild type lacking an epiallele (RS or RW). In contrast, paramutation occurs in RRSS (tetraploid) hybrids and is revealed by a reduced number of resistant F2 plants, significantly lower than in the control crosses of R with W (RRWW), independent of whether the S allele came from the maternal or paternal side, but with variation between F2 populations from different F1 parents [17]. This variation raised the question whether external factors during the growth of the F1 hybrids could influence the penetrance of paramutation. We hypothesized that temperature is an important parameter, because its influence on a paramutation-resembling inheritance in Drosophila and the expression of the paramutable *r* gene in maize had been reported [23, 24]. We grew diploid and tetraploid F1 plants obtained from crosses between R and S (or W) at 10°C, 19°C, or 24°C during their flowering and seed set period (Fig 2A). We germinated F2 seeds on hygromycin selection medium and scored the ratio of resistant seedlings (Fig 2B and 2C). F2 from tetraploid RRSS hybrids grown at 19°C (the standard growth conditions) confirmed the results from earlier experiments, with a reduced ratio of resistant plantlets compared to the RRWW controls. This difference was even more significant in the progeny from parents grown at 24°C, indicating an enhanced interaction between R and S. Unexpectedly, progeny from RRSS and RRWW plants grown at 10°C showed no difference in the resistance assay, indicating that the presence of the S epiallele did not lead to paramutation of R at the lower temperature (Fig 2B). All F2 plants from diploid hybrids contained approximately 75% resistant seedlings, excluding any influence of the S epiallele under all tested temperature conditions (Fig 2C). Therefore, paramutation occurs at 19°C but not in plants grown at 10°C. These results confirm that paramutation in this system is associated with tetraploidy and indicate that its penetrance is indeed affected by external factors, exemplified here by temperature.

**Figure 2.**
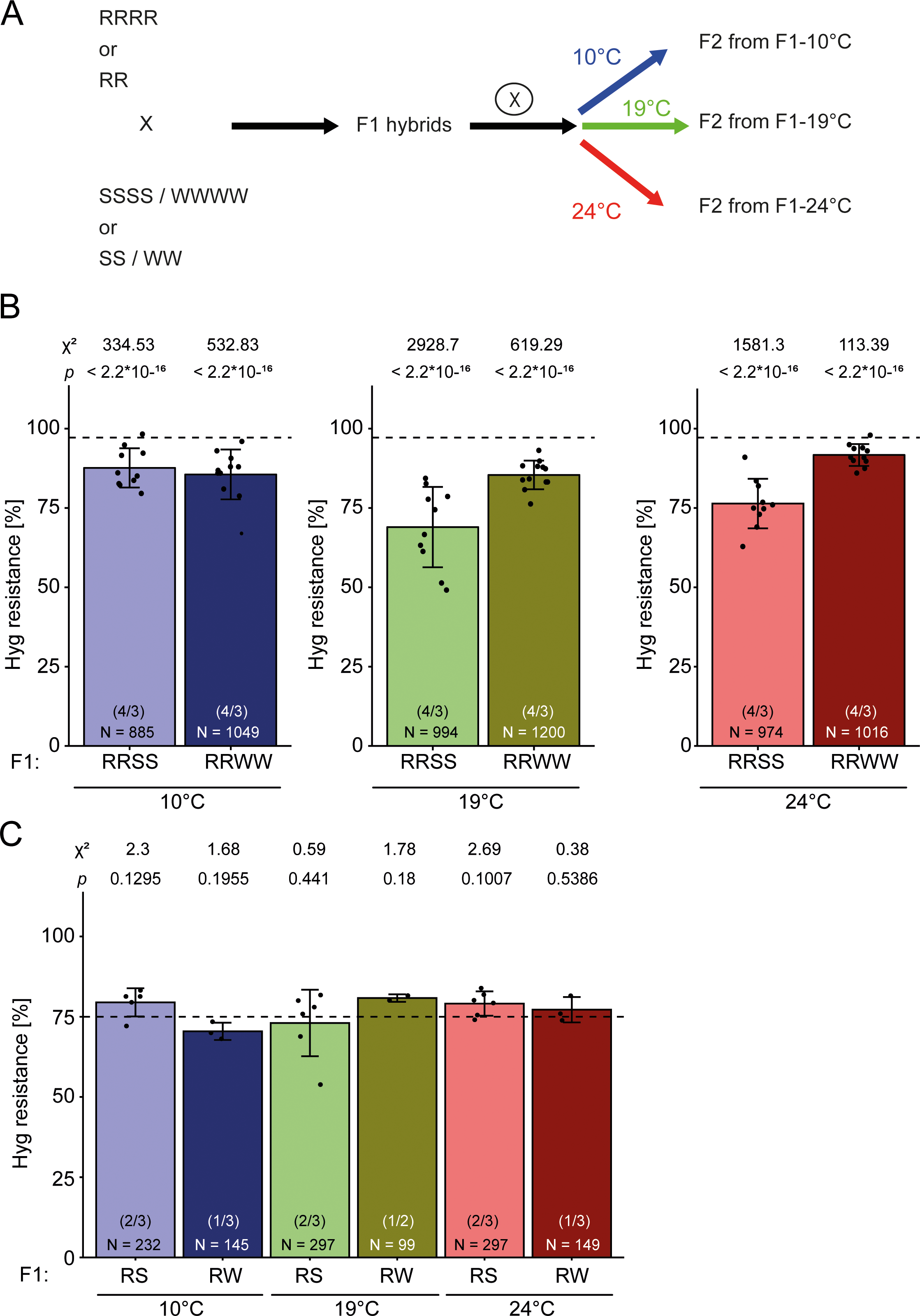
The degree of paramutation in tetraploids depends on growth temperature in F1. (A) Experimental setup: Diploid or tetraploid plants with R epialleles were crossed with those containing S or W, and F1 hybrids from reciprocal crossings were grown at 19°C for three weeks before transfer to either 10°C, 19°C, or 24°C until seed maturity. F2 seeds were germinated at 19°C on GM plates containing 20 mg/L hygromycin B and resistance ratios determined after 14 days for tetraploids (B) and diploids (C). Data for reciprocal crosses were combined, as no parent-of-origin difference was observed. Number in parentheses: different F2 populations / technical repetitions for each population. N = number of tested seedlings in each group. Bars represent the mean from two biological with three technical replicates each (B) and one biological with three technical replicates (C), with 100 plated seeds (B) each, or 50 plated seeds (C). Error bars indicate standard deviation. Dashed lines represent the expected segregation for tetraploid (B, 35:1 ≙ 97.2%) and diploid F2 populations (C, 3:1 ≙ 75%). F1 growth temperature is indicated by colour: 10°C (blue), 19°C (green), 24°C (red); light colours: F2 of R/S hybrids, dark colours: F2 of R/W hybrids. Statistical analysis was performed by summed Chi-square goodness-of-fit test with the indicated values.

### Epialleles involved in Arabidopsis paramutation have different sRNA signatures

Multiple lines of evidence indicate that *trans*-silencing phenomena, such as paramutation, involve the production of small RNAs (sRNA) that silence the paramutated allele. The presence of sRNAs at loci participating in paramutation has been shown for genes in maize [25, 26], tomato [9], and Arabidopsis [27], including a very limited analysis of the S allele in this study [20]. To address the possible mechanistic role of sRNA, we generated sRNA libraries from total RNA of 14 d-old seedlings of tetraploid and diploid homozygous R and S lines and mapped reads with a length from 18-26 nt to the Arabidopsis TAIR10 reference genome extended *in silico* with the inserted *HPT* epiallele. Whereas the size distribution of sRNAs mapping to the whole genome were similar in all libraries, there was a striking difference regarding the size classes between lines carrying the different epialleles: we found mainly 21 nt sRNAs at R and mostly 24 nt sRNAs at S, regardless of ploidy (tetraploids: Fig 3A, diploids: S1A Fig). Both size classes of sRNAs mapped predominantly to the duplicated region (promoter and vector sequence) of the epiallele (Fig 3B, S1B Fig), indicating that these sRNAs may originate from either copy, although their origin from the downstream repeat is more likely as this is part of the incompletely terminated transcript [20]. Libraries prepared from wild type plants without the reporter contain insignificant low numbers of reads mapping to the reporter (S2B Fig), confirming the origin of the 21/24 nt reads in Fig 3B from the epialleles. The abundance of the 24 nt sRNAs was similar between diploid and tetraploid S lines, and they were distributed along the whole duplicated promoter/vector region. In contrast, tetraploid R lines had higher levels of 21 nt sRNAs than diploids. Many of these reads mapped to a specific region of the promoter. The *HPT*-ORF and the non-coding region downstream of the second promoter copy were almost devoid of sRNAs in libraries from all samples (Fig 3B, S1B Fig). These data indicate that the active and silent epialleles are characterized by matching sRNAs of different length, with specific profiles and quantities.

**Figure 3.**
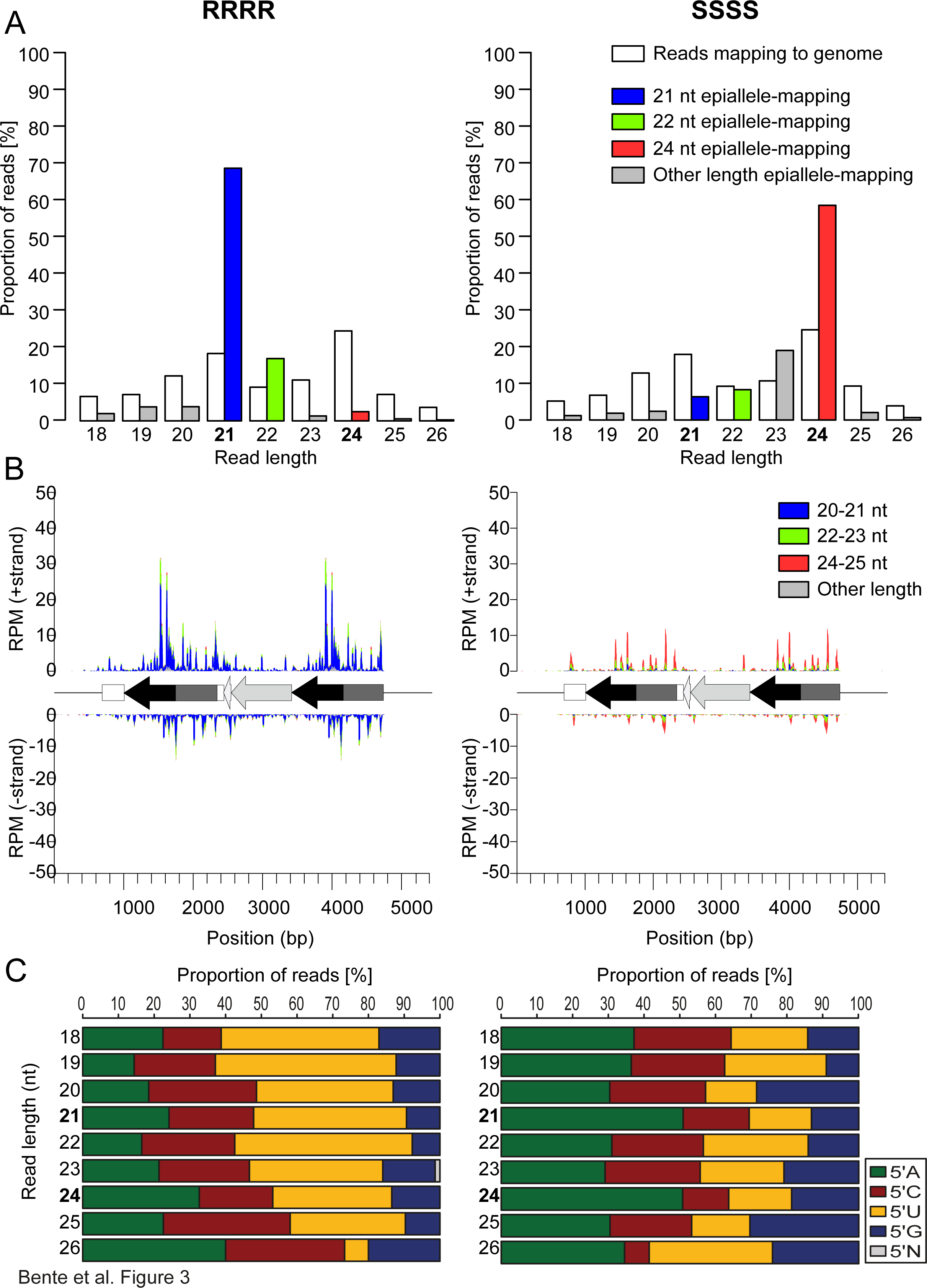
Epialleles are associated with distinct classes of sRNAs. (A) Proportion of sRNA length from all reads mapped to the epialleles in 14 day-old seedlings with tetraploid active (RRRR) or silenced (SSSS) epialleles. See S1 Fig for the corresponding data from diploids. (B) Coverage plots of 18 – 26 nt sRNA along the RRRR or SSSS epialleles. (C) Proportion of 5’ prime nucleotides of epiallele-specific sRNAs in RRRR (left) and SSSS (right) plotted by size. See S1 Fig for the corresponding data for reads mapped to the genome.

The frequency of the four bases at the 5’end of sRNAs correlates with their length and serves as an indicator for their potential association with specific Argonaute (AGO) proteins [28]. In all lines, sRNAs mapping to the wild-type genome confirmed the known preference for a 5’U in 21 nt sRNAs and a 5’A in 24 nt sRNAs (S2A Fig). The 5’ base distribution of epiallelic sRNAs in both tetraploids (Fig 3C) and diploids (S1C Fig) matches this preference: 5’ nucleotides of the 21 nt sRNAs mapping to the R epiallele were biased for 5’U, indicating a preference for AGO1, while half of the 24 nt sRNAs from the S epiallele had an A at the 5’ end, suggesting association with either AGO4 [28] or AGO3 [29].

### sRNA signature and mutant analysis support involvement of RdDM

The relatively high percentage of 5’A in the 24 nt sRNAs from the S epiallele (Fig 3C and S1C Fig) suggested the involvement of RNA-directed DNA methylation (RdDM) [30, 31], as suggested by several mutational studies in maize paramutation [32]. The limitation of the R/S interaction to tetraploid plants made any forward-directed screen impractical, as recovery of recessive mutations could only be expected in M3 generations and with extremely low probability. However, the efficiency of CRISPR-Cas9- based mutagenesis in Arabidopsis [33] allowed us to test the hypothesis that RdDM is involved in this case of paramutation. Paramutation between the *B-I* and *B’* allele in maize requires functional RDR2 [11, 34] and several subunits of the plant-specific DNA-dependent RNA polymerases IV and V [35–38]. Therefore, we targeted two corresponding Arabidopsis genes, *RDR2* (At4g11130) and the largest subunit of PolIV (*NRPD1*, At1g63020) in tetraploid R and S plants for CRISPR-Cas9- based mutagenesis. We selected lines homozygous for the desired mutations and the absence of the Cas9 gene by genotyping. Loss of NRPD1 function was demonstrated by the genome-wide absence of endogenous 24 nt small interfering RNAs (S3A Fig). Furthermore, the mutation in *NRPD1* led to depletion of epiallele-derived 24 nt sRNAs in SSSS (S3B Fig). Unlike the *NRPD1* mutant *rmr6* in maize, which reactivated the silent state of *r1* and *pl1* alleles [36], the newly generated Arabidopsis *cas9nrpd1* mutant did not lead to reactivation of the S epiallele. We tested the potential for paramutation by generating F1 hybrids homozygous for the mutation and scoring F2 seedlings for hygromycin resistance, together with progeny from corresponding wild type combinations grown in parallel. The ratio of resistant to sensitive progeny from mutant RRSS hybrids was much closer to that of the RRWW crosses (Fig 4A) than in the wild-type background (Fig 4B). This indicates a reduced establishment of paramutation when RdDM is impaired, but that RdDM does not play a role in the maintenance of the silent state of the S epiallele.

**Figure 4.**
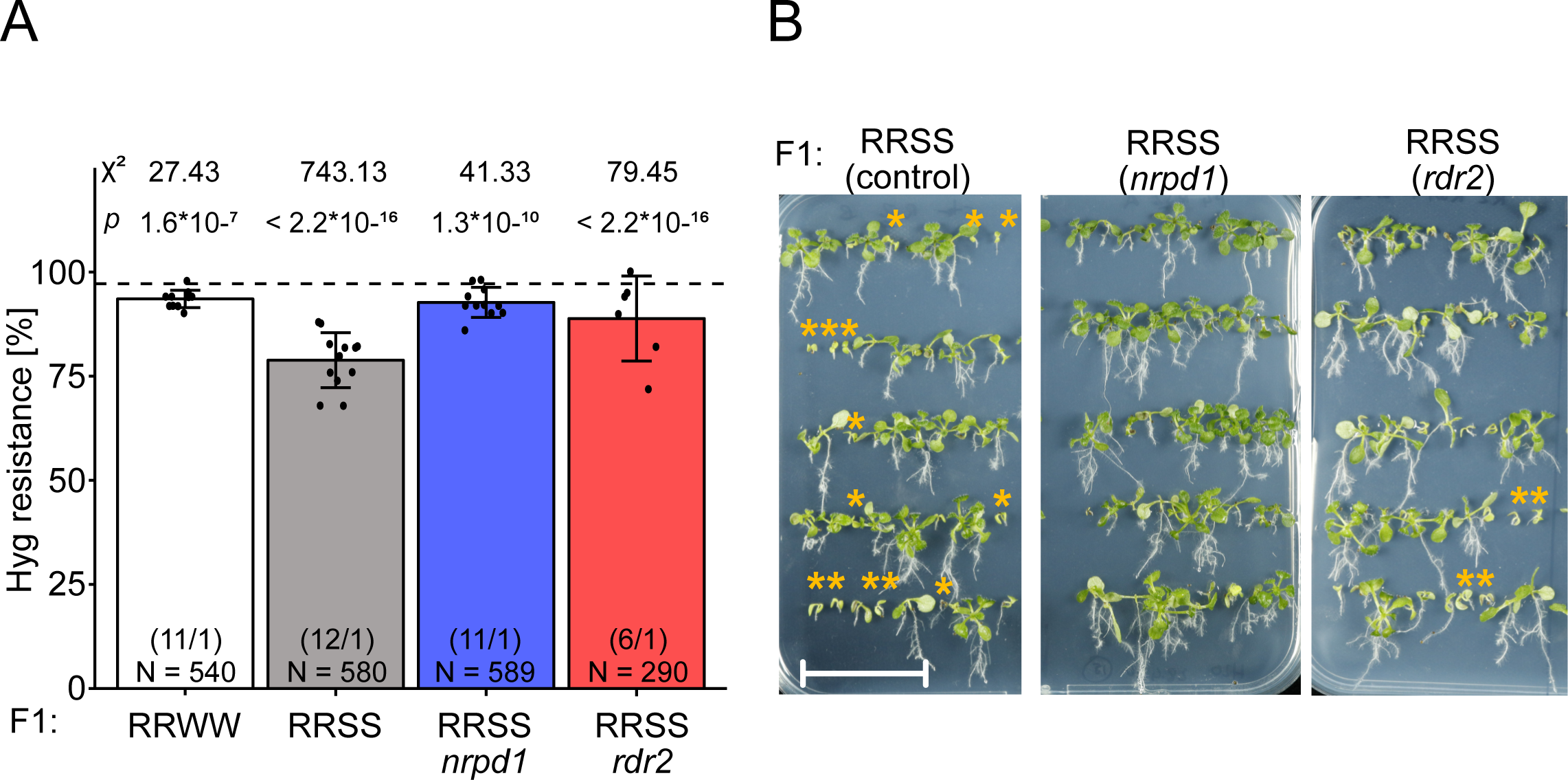
Lack of paramutation in the background of RdDM mutants. (A) F2 seeds obtained by selfing tetraploid F1 hybrids were germinated on GM plates containing 20 mg/L hygromycin B and resistance ratios determined after 14 days. RRWW control (white), paramutation test hybrid RRSS in wild type (grey), *nrpd1* mutant (blue) or *rdr2* mutant (red). Data for reciprocal crosses were combined. Number in parentheses: different F2 populations / technical repetitions for each population. N = number of tested seedlings in each group. The dashed line represents the expected F2 segregation for tetraploids (97.2%), statistical analysis is based on a summed Chi-square goodness-of-fit test with indicated values. (B) Representative images for resistance assays quantified in (A). Orange asterisks indicate sensitive seedlings, scale bar = 3 cm.

### The active epiallele is sensitive to silencing by *trans*-acting small RNAs

The results of the mutant analysis suggested that sRNAs are important for establishing paramutation. We therefore tested the hypothesis that sRNAs were sufficient to silence the otherwise stable R epiallele. We designed three hairpin (hp) constructs that would induce strong sRNA formation targeting different regions within the duplicated vector/promoter region (hp1/2/3; Fig 1A) and transformed RR and RRRR plants. Seeds from individual T1 plants were sorted for the presence or absence of the hairpin construct via seed-specific GFP expression, which was encoded in the vector, and northern blot analysis documented processing of the transcript into small RNAs (S4 FigB). Plants were grown with or without hygromycin. Under selection, all hairpin variants caused reduced root growth in seedlings, compared to plants without hairpins (Fig 5A and 5B), indicating silencing of the R allele in diploid and tetraploid lines. The degree of silencing was highest with hp3 which is homologous to the promoter. However, hp1 and hp2 which are homologous to the vector sequence further away from the promoter within the epiallele also resulted in significant growth inhibition in seedlings under selection (Fig 5A). To quantify the effect of the hairpins on transcript levels at a later stage of development, we determined *HPT* expression by qRT-PCR in flower buds from diploid and tetraploid T1 plants with and without hairpins sorted as above. In diploid plants, all three hairpin variants reduced *HPT* expression to a similar level (Fig 5C left panel), whereas the effect increases with proximity of the hp-matching region to the transcription start site of the *HPT* gene in tetraploid plants (Fig 5C right panel). The expression levels of the hairpin constructs are expected to be averaged as we sampled many T1 plants representing independent insertion sites. Therefore, the hairpin constructs appear to silence the two R copies in diploids more efficiently than the four R copies in tetraploids. These data indicate that stoichiometry between sRNAs and the target genes could explain the role of ploidy in the response to silencing cues.

**Figure 5.**
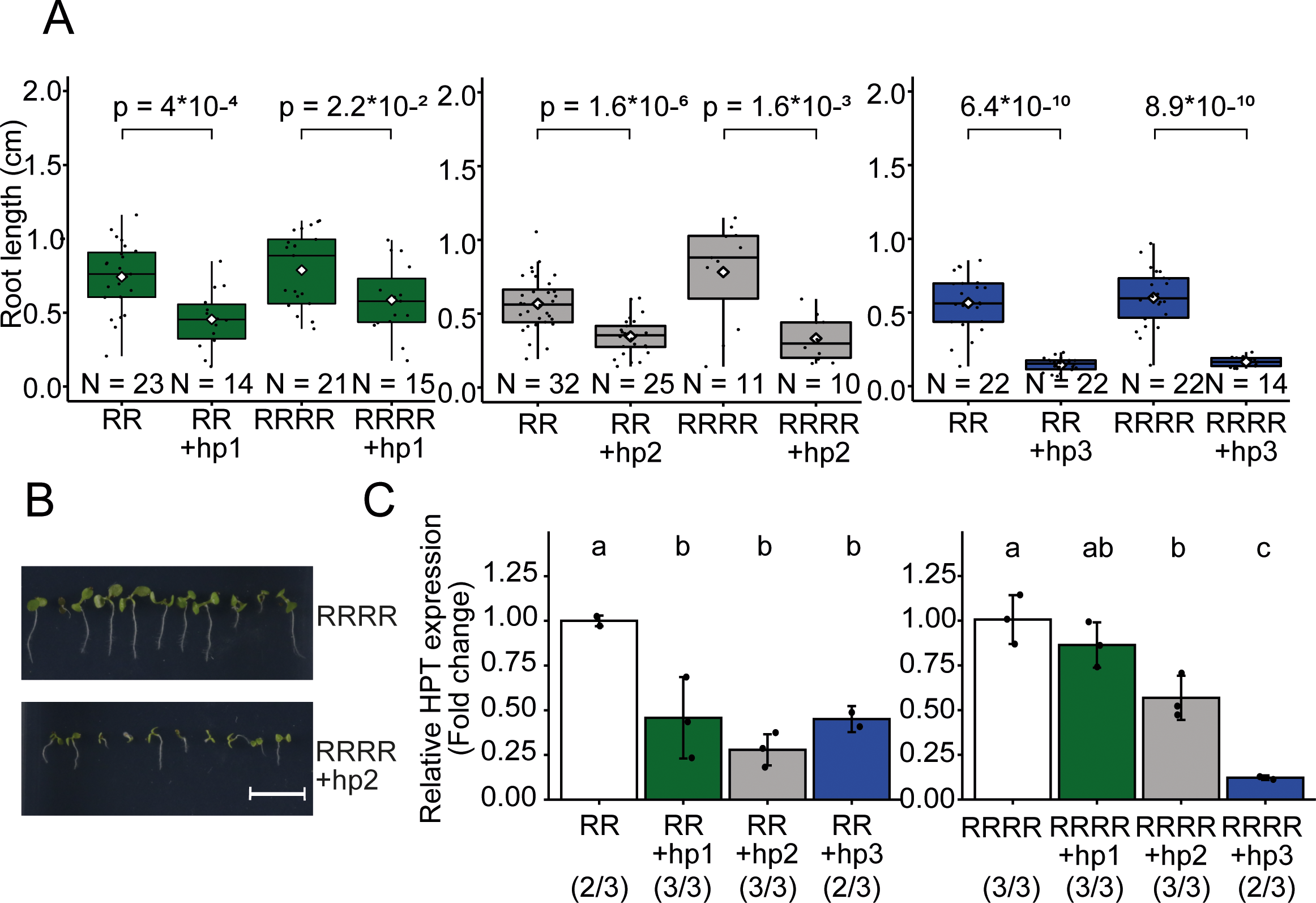
Ectopic RNAi expression reduces *HPT* expression to different extent in diploids and tetraploids. (A) Three constructs with constitutive expression of hairpin-forming palindromes homologous to either the vector region of the R epiallele (hp1, green; hp2, grey) or the promoter region (hp3, blue) were inserted in ectopic positions, and their transcripts are processed into sRNAs. (A) Box plots of root length (measured with SmartRoot software) as indicator of resistance in the presence of 20 mg/L hygromycin in 7 d-old T1 seedlings grown from seeds of T0 plants with or without the hairpin construct (sorted by seed-specific GFP fluorescence provided by the construct and grown on the same plate). White diamonds indicate the mean of each group. Statistical analysis was made by Welch’s two sample T-test, p-values of relevant comparisons are indicated. N = number of tested seedlings in each group. (B) Representative images for root growth in lines without (upper panel) and with (lower panel) the hairpin construct, scale bar = 1 cm. (C) Relative *HPT* expression in flower bud tissue of hairpin-containing T1 plants in tetraploids (left panel) and diploids (right panel), compared to sibling plants without hairpins. Data were normalized to the housekeeping gene *SAND (At2g28390)* and shown as fold-change values. Number of biological and technical replicates are indicated in parentheses. Bars indicate standard deviation of biological replicates. Statistical analysis was performed by one-way ANOVA followed by a post-hoc Tuckey HSD test (α = 0.05).

### New silenced epialleles can exert secondary paramutation

The epialleles do not contain an inverted repeat that could form hairpins to serve as a source of sRNAs. However, tandem-oriented repeats can affect paramutation, depending on their copy number [39–41]. Evidence for a role of the direct repeat of the vector/promoter region in our paramutation system came from the previously described structural variants (RΔ, derived from the S epiallele by mutagenesis), in which rearrangements downstream of the *HPT*-ORF led to reactivation of the upstream promoter [20]. We made use of two different, independent deletions to ask whether and how they would affect the paramutability. In RΔ2, the whole transgenic sequence downstream of the defective terminator (including the vector/promoter repeat) was deleted, whereas RΔ4 lacked only the terminal non-coding carrier sequence, maintaining the repeat (Fig 1A). Tetraploid, homozygous derivatives of both lines (RΔ2RΔ2RΔ2RΔ2 and RΔ4RΔ4RΔ4RΔ4, selected after colchicine treatment) were crossed with SSSS or WWWW lines (Fig. 1B) and the F2 progenies screened for hygromycin resistance. Only F2 offspring from reciprocal crosses between RΔ4RΔ4RΔ4RΔ4 x SSSS (RΔ4RΔ4SS), but not from RΔ2RΔ2RΔ2RΔ2 x SSSS (RΔ2RΔ2SS) crosses, showed reduced hygromycin resistance like crosses with full length RRRR (Fig 6A). This indicates that RΔ2 is a neutral allele, whereas RΔ4 is still paramutable.

**Figure 6.**
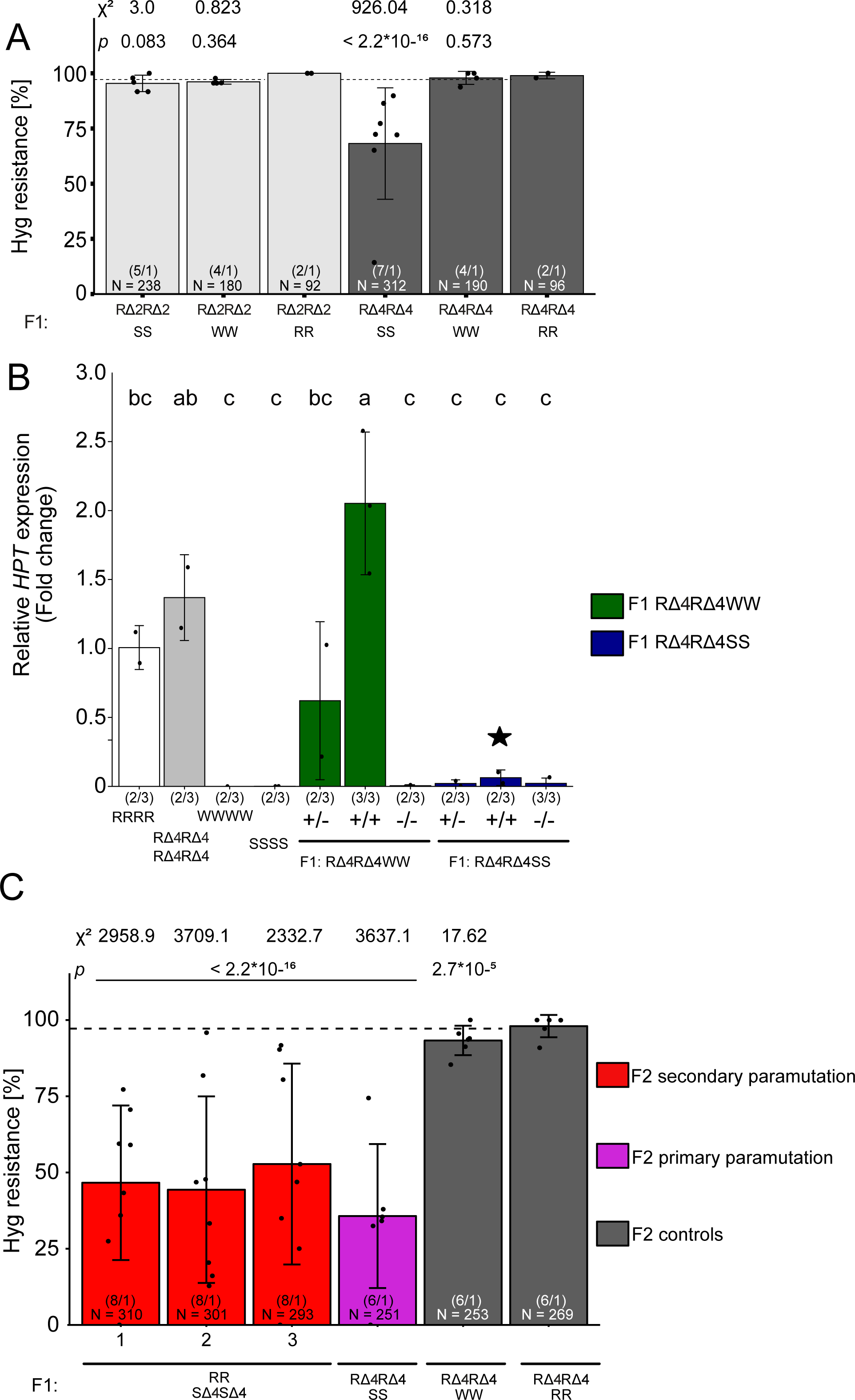
Paramutation depends on epiallele structure, and paramutated epialleles can exert secondary paramutation. (A) Tetraploid F2 seeds obtained by selfing F1 hybrids of crosses between S/W/R and RΔ2 (light grey) or RΔ4 (dark grey) were germinated on GM plates containing 20 mg/L hygromycin B and resistance ratios determined after 14 days. Data for reciprocal crosses were combined. Number in parentheses: different F2 populations / technical repetitions for each population. N = number of tested seedlings in each group. The dashed line represents the expected F2 segregation for tetraploids (97.2%), statistical analysis is based on a summed Chi-square goodness-of-fit test with indicated values. (B) RΔ4 deletion alleles were combined with either W or S partners (all tetraploid). Individual plants in F2 progeny were genotyped for RΔ4, sorted into groups heterozygous (+/-) or homozygous (+/+) 27 for the deletion and material from14 d-old seedlings subjected to *HPT* expression analysis relative to homozygous RRRR and the housekeeping gene *EIF4A1 (At3g13920)*. Number in parentheses: biological / technical replicates. Error bars indicate standard deviation of biological replicates. Statistical analysis was performed by one-way ANOVA followed by a post-hoc Tuckey HSD test (α = 0.05). The black star indicates individuals homozygous for the R allele with the deletion and lacking any S copy, nevertheless with substantially reduced HPT expression. (C) Plants from primary paramutation crosses, homozygous for RΔ4 alleles (excluding the presence of the old S allele) and only residual *HPT* expression (black star in B, now called SΔ4SΔ4SΔ4SΔ4, Fig. 1) were crossed to plants with the active, full length R alleles and analysed for hygromycin as described before in F2, in parallel to controls. The dashed line represents expected F2 segregation for tetraploid populations at 97.2%; statistical analysis is based on a summed Chi-square goodness-of-fit test with indicated values. N = number of tested seedlings in each group. Number in brackets: different F2 populations / technical repetitions of each population.

The most persuasive evidence for paramutation is acquired paramutagenicity of a paramutated allele, i.e. the ability to exert secondary paramutation. In the initial paramutation cross RRRR x SSSS, paramutated R alleles do not have any sequence polymorphism that distinguishes them from S alleles in the segregating F2. To enable identification of the paramutated allele after its conversion, we used the deletion in RΔ4 for PCR-based genotyping in the progeny of RΔ4RΔ4SS plants. We selected individual F2 plants that were either homozygous (+/+) or heterozygous (+/-) for the deletion and quantified *HPT* expression with qRT-PCR. As expected from previous studies and in accordance with the hygromycin resistance assay, RRRR and RΔ4RΔ4RΔ4RΔ4 showed strong expression, while there were no detectable transcripts in SSSS and WWWW. Moreover, only progeny from RΔ4RΔ4WW hybrid parents expressed *HPT* in comparable amounts (Fig 6B). In contrast, progeny from RΔ4RΔ4SS parents had very low *HPT* expression, including plants homozygous for the previously active RΔ4 allele. These plants contained only paramutated copies (Fig 6B, black star), indicating strong suppression by the S allele in the previous generation lasting after segregation. Therefore, all RΔ4 alleles had undergone paramutation to SΔ4SΔ4SΔ4SΔ4 (Fig 1B). We used these plants to assay secondary paramutation. Tetraploid hybrids from crosses between plants with the fully paramutated alleles (SΔ4SΔ4SΔ4SΔ4) and plants with the initial active R alleles (RRRR) (Fig 1B) resulted in F2 populations with a similarly reduced ratio of hygromycin resistance as obtained after the primary paramutation (Fig 6C). This indicates that once an *HPT* allele is paramutated, it can reduce *HPT* expression from a naïve R allele in the process of secondary paramutation.

### Removal of the downstream duplication does not reactivate a silent upstream promoter but boosts expression from active epialleles

RΔ2 and RΔ4 alleles had been identified in a forward genetic screen selecting for reactivation of the S epiallele [20]. Both deletions resulted in hygromycin resistance, but RΔ2 lost paramutagenicity while RΔ4 maintained this feature. The different, randomly induced structural changes (Fig 1A) suggested that the removal of the duplicated downstream region in RΔ2, but not in RΔ4, might explain the difference. To precisely recapitulate this event, we applied CRISPR gene editing to eliminate the downstream repeat as well as the non-coding sequence from R and S alleles. Cas9-encoding transgenes with suitable sgRNAs were transformed into diploid and tetraploid plants. T1 plants were selfed or outcrossed and the progeny genotyped for homozygosity of the wanted deletion and the absence of the Cas9 transgene. PCR amplification across the deletion revealed the same amplicon size, and Sanger-sequencing of the amplicons of two randomly chosen lines revealed the same ligation between cuts 3 bp upstream of the respective PAM sites. To distinguish the new deletions from the previously described alleles, the Cas-9 induced alleles are denoted as Rγ and Sγ (Fig 1A).

We screened diploid (RγRγ, SγSγ) and tetraploid (RγRγRγRγ, SγSγSγSγ) plants for hygromycin resistance. R-derived plants with the deletion grew longer roots on hygromycin (Fig 7A and 7B). Northern blots with RNA from these lines confirmed a shorter transcript of ∼1.6 kb (as excepted for the deletion), with ∼3-4x more signal strength than in R plants, in both diploids and tetraploids (Fig 7C). These data indicate that, despite high transcript levels and stable hygromycin resistance from full length R epialleles, the associated 21 nt RNAs might contribute to partial reduction of transcripts originating from the upstream promoter.

**Figure 7.**
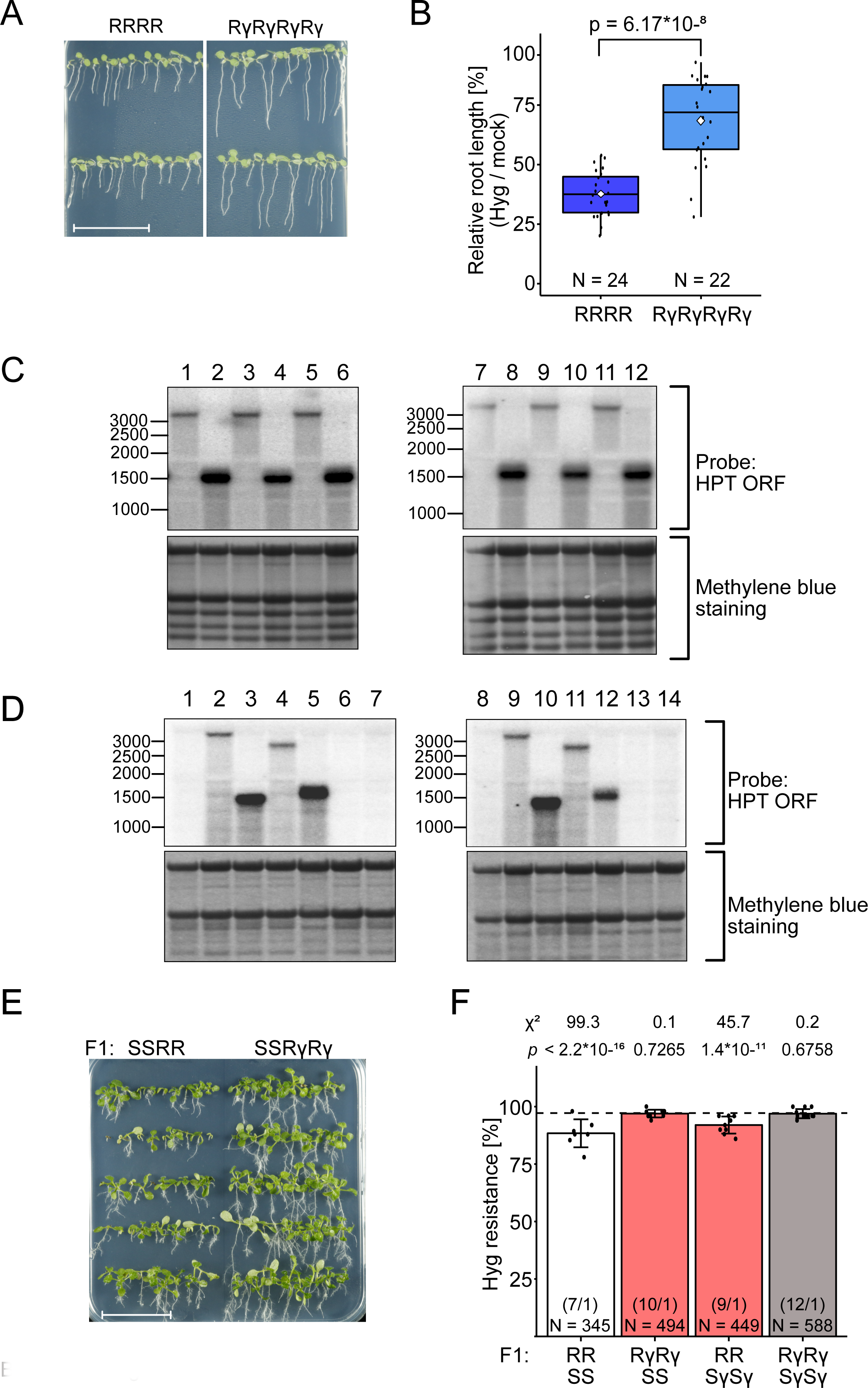
Deletion of the downstream repeated region leads to higher expression at active epialleles and reduces paramutation. (A) Representative images of root growth assays in the presence of 20 mg/L hygromycin with 7 d-old plants containing full length or truncated active epialleles. Scale bar = 3 cm. (B) Box plots of relative root length on hygromycin versus non-selective medium as in (A) (measured with SmartRoot software). White diamonds indicate the mean of each group. Statistical analysis was performed with Welch’s two sample T-test with indicated p-value. N = number of tested seedlings in each group. (C) Northern blots of *HPT* transcripts in seedlings of independent lines of tetraploid (1-6) and diploid (7-12) plants with full length (1,3,5,7,9,11) or truncated (2,4,6,8,10,12) active epialleles. (D) Northern blots of *HPT* transcripts in flower buds of tetraploid (1-7) and diploid (8-14) plants. 1: WWWW; 2: RRRR; 3: RΔ2RΔ2RΔ2RΔ2; 4: RΔ4RΔ4RΔ4RΔ4; 5: RγRγRγRγ; 6: SSSS; 7: SγSγSγSγ; 8: WW; 9: RR; 10: RΔ2RΔ2; 11: RΔ4RΔ4; 12: RγRγ; 13: SS; 14: SγSγ. Location of the probe for blots in C and D is illustrated in Fig. 1. (E) Representative image for hygromycin resistance assays with 14 d-old F2 plants after crossing parents with full length or truncated epialleles. Scale bar = 3 cm. (F) Resistance ratios calculated from assays like in (E). Data from reciprocal crosses were combined, statistical analysis is based on a summed Chi-square goodness-of-fit test with indicated values. Number in parentheses: different F2 populations / technical repetitions for each population. N = number of tested seedlings in each group.

In contrast to the deletions after random mutagenesis [20], S-derived plants with the targeted deletions (SγSγ, SγSγSγSγ) remained completely sensitive and without detectable *HPT* transcripts (Fig 7D). Therefore, the removal of the duplicated regions was not sufficient to restore the *HPT* expression. The sRNA profiles of the deletion lines were quite similar between diploid and tetraploid lines, and between S and Sγ (S5 Fig B and D). Unexpectedly, the profiles of both *HPT*-expressing deletion alleles (RΔ2, Rγ) lost the 21 nt sRNAs characteristic for the full-length R allele but gained 24 nt sRNAs along the remaining vector- promoter region (S5 C and F). These results demonstrate that removal of the downstream repeat eliminates the 21 nt class and further implies that the occurrence of 24 nt sRNA does not *per se* lead to silencing. The main difference in the profiles of active (RΔ2, Rγ) versus silent (Sγ) deletion alleles is the presence of sRNAs along the *HPT* coding region in the latter (S5 Fig C, D, and F). Those at the vector-promoter regions are obviously not interfering with expression and might be remnants from the previously present duplication, resembling the peaks of 24 nt sRNAs at the tandemly orientated LTR regions of transposons in several plants [42–45].

To determine if Sγ and Rγ could participate in paramutation, we performed reciprocal crosses between tetraploids with full length or deleted R and S alleles and screened F2 segregation under hygromycin selection. Resistance in progeny from RRSγSγ hybrids was slightly higher than for RRSS parents (Fig 7E and 7F), indicating that the deletion of the repeat might affect paramutagenicity. RγRγSS did not longer show paramutation, probably due to higher HPT transcript level from the shortened R allele (Fig 7E and 7F). Taken together, the removal of the duplicate and the absence of 21 nt sRNAs boosts expression from the active epiallele and renders it less or not at all sensitive to paramutation, while the removal of the duplication from the silent epiallele weakens its paramutagenicity. This implies that the structure of the epialleles on both partners, in combination with the specific small RNA profiles, influence the degree of paramutation.

## DISCUSSION

In this study, we investigated a case of paramutation in *Arabidopsis thaliana* that shares the main features, including secondary paramutation, with other examples, while being different with regard to its timing and link to polyploidy. We documented important roles for sRNAs to initiate silencing, the structure of the allele itself, the environmental conditions during growth of the hybrids, the developmental stage, and the copy number ratio between the epialleles. We describe a model that makes these features plausible (Fig 8): the potentially strong *HPT* expression from the upstream Cauliflower Mosaic Virus 35S promoter is weakened in the active epiallele by 21 nt sRNAs connected with the downstream repeat, but still remains strong enough to produce sufficient HPT protein to detoxify hygromycin. Regular F2 resistance segregation after crosses with the “empty” wild type imply that one active allele, even with the repeat, is enough to render a plant resistant, regardless of whether the dosage is 1 out of 2 (diploid) or 1 out of 4 (tetraploid). If the cross introduced the inactive epiallele, *HPT* expression is still maintained high enough to allow resistance as long as the associated 24 nt sRNA are present in limited amounts. The ectopic overexpression of sRNA varieties from the hairpin constructs demonstrated that they are triggers of rapid, efficient, and lasting trans-silencing, as indicated by reduced HPT transcript in T2 progeny devoid of the hairpin construct (S4 FigC). However, in R and S lines without hairpin constructs, 21 nt and 24 nt sRNA, respectively, are not detectable on northern blots, even after sRNA enrichment or with sensitive LNA probes, and can only be quantified by their relative contribution of reads in the sRNA libraries. A comparison of absolute amounts is therefore difficult. However, in diploid hybrids, both epialleles are present in a 1:1 ratio, which likely does not tilt the balance towards silencing. In tetraploids, 22.2% of the F2 progeny will contain one active and three silent copies; here, the abundance of 24 nt sRNAs cross a threshold for initiating the trans-silencing effect. The “missing” portion of resistant plants in paramutation cross progeny is usually between 15-30%, compatible with the assumption that especially plants with more silent than active epialleles (never occurring in diploids) are undergoing the switch. This is further supported by a limited analysis of progeny from a backcross between RRSS hybrids with SSSS, expected to have 1 R and 3 S copies in 2/3 of the population, which was completely hygromycin-sensitive (S6 Fig). A similar higher incidence for paramutation with a triplex genotype for the paramutagenic *sulfurea* allele is apparent in tetraploid tomato hybrids [46], and paramutation can occur independently at single alleles in subsequent generations [47].

**Figure 8.**
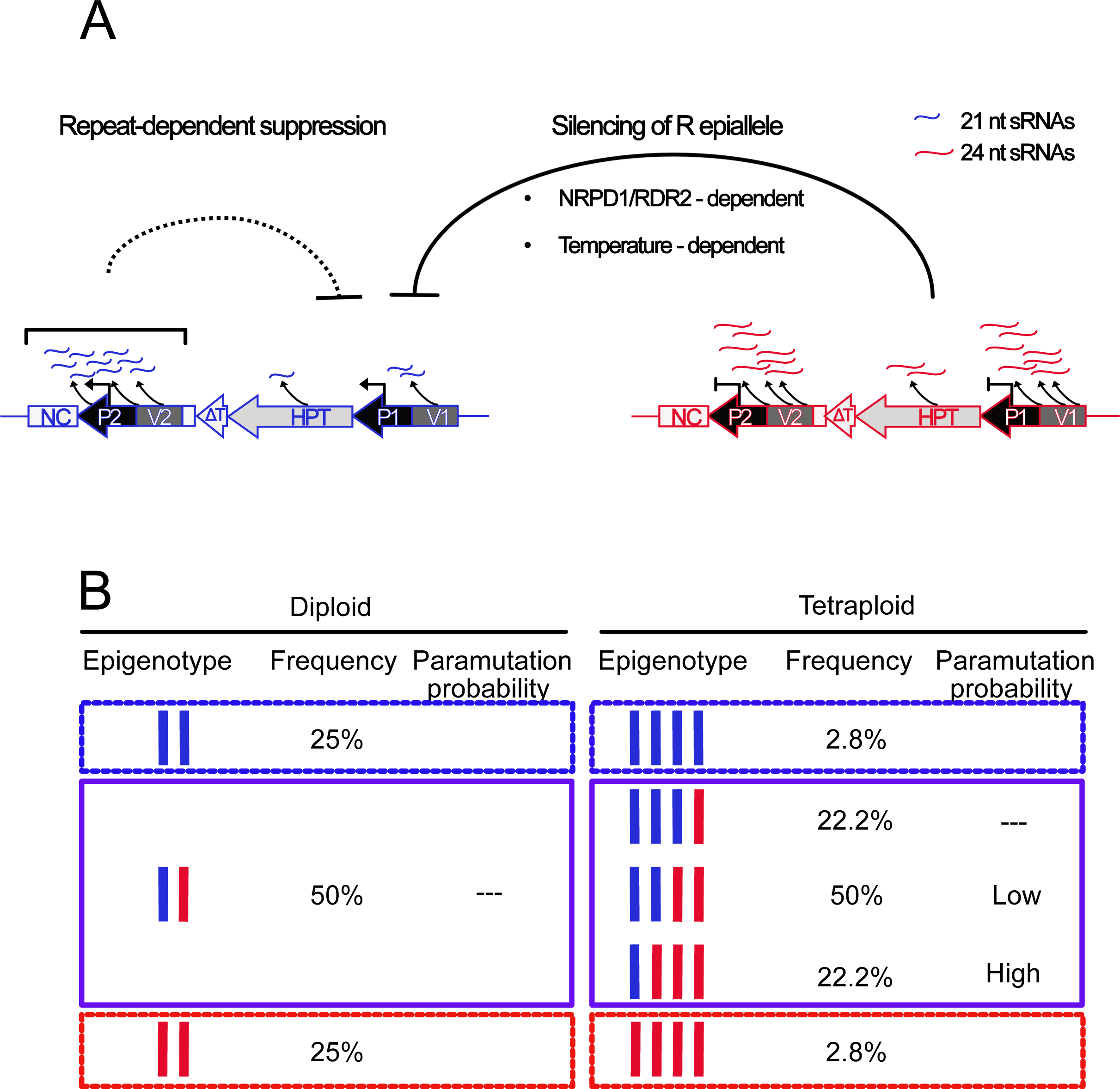
Features of polyploidy-dependent paramutation. (A) The expressed epiallele (blue) confers hygromycin resistance, although the downstream repeat and the 21 nt sRNA limit the amount of transcripts from the upstream promoter to some extent. The silent epiallele (red) is associated with 24 nt sRNAs and exerts strong silencing *in trans*, involving temperature-dependent RdDM. (B) Distribution of epigenotypes in F2 populations.Paramutation occurs in a dosage-dependent way and may be restricted to hybrids with more silent than active copies. This combination does not occur in diploids and explains the limitation of paramutation to tetraploids.

The Arabidopsis/*HPT* and the tomato/*sulf* system are the only cases for which a role of tetraploidy in paramutation has been described. In neither case is the silent state of the paramutagenic epiallele connected with polyploidy, as both alleles are stable in diploid background, but paramutation seems to be supported by the 4x karyotype. Genome duplications after formation of polyploids has often been associated with epigenetic changes [48–52] and might trigger the origin of epialleles. However, most studies were performed with allopolyploids, in which combining different genomes modifies chromatin, DNA methylation, or small RNA profiles. In these cases, changes can be due to either increased chromosome numbers, increased genetic diversity, increased gene copy numbers, incompatibility of regulatory components, or a mixture of these parameters. The situation in autopolyploids is simpler, as it excludes genetic determinants. Newly created autopolyploids can contain epigenetic modifications, e.g. switchgrass [53] and Arabidopsis [54], but effects on gene expression are much smaller than in allopolyploids [55] and occur rather stochastically [56]. We did not find prominent differences between diploid and tetraploid lines, besides the divergent *HPT* expression and small RNA profiles at the epialleles. Therefore, the limitation of paramutation to the tetraploid hybrids is likely not due to the chromosome number as such, or to indirect consequences like larger nuclei [57], cell cycle regulation [58], or differences in meiosis [59]. It is more plausible to assume that rather the copy number of the epialleles, or better their ratio, determine the epigenetic change. Copy numbers, not varied by ploidy but by allelic variation of repetitive sequences, are also a major determinant of paramutation in maize [39–41, 60], again mostly in tandem configuration but sometimes far upstream of the coding sequence.

Another paradigm for a dosage effect is transcriptional silencing of a transposon, which occurs only after the copy number of the genomic source has increased beyond a threshold by new insertions [61]. A striking similarity with *HPT* paramutation is the role of 24 nt sRNAs that characterize the paramutagenic epiallele and that appear in connection with the switch to transcriptional transposon silencing. 24 nt sRNAs were also reported in very low amounts for the *B-I*/*B’* paramutation in maize, depending on the RDR2 ortholog MOP1, but were also found in neutral alleles and therefore not sufficient for paramutation [62]. As in our study, their ectopic expression in larger amounts can mimic paramutation [62], again supporting a role for dosage dependency. There is no report of 21 nt sRNAs at paramutable maize alleles. This is in contrast to the *HPT* system, where their presence might be connected to the downstream location of the repeat and its presence in the transcript. However, the repeats are in tandem orientation, so that hairpin formation as a precursor for double-stranded RNA is unlikely. The requirement for MOP1 in maize paramutation [11] and the reduced paramutation in *rdr2* mutants in our study strongly support a dependence on RNA-dependent RNA polymerase for the creation of the 24 nt small RNAs. As MOP2 and MOP3 are also components of RdDM-like silencing [37, 63], it is clear that this epigenetic regulatory pathway is a core component of paramutation. Even outside of paramutation siRNA-induced epigenetic changes can persist over several generations [64]. However, the interference with RdDM in the Arabidopsis system did not result in hygromycin resistance, in contrast to the derepression of both *B’* and *P’* leading to increased pigmentation e.g. in *mop1* [11]. However, this could be due to the exceptionally strong silencing at the S epiallele, which needs simultaneous removal of DNA methylation and repressive histone modifications [18].

The role of the 21 nt sRNAs in the Arabidopsis system remains elusive. They are only present together with the full-length active R allele regardless of ploidy and clearly dependent on the presence of the downstream repeat, as they disappear after its removal in Δ and γ deletion lines. 21 nt RNAs are usually associated with post-transcriptional silencing (PTGS) [65], characterized by the 5’ preference for U and associated with AGO1 [28] and guide mRNA cleavage or translational repression [66]. The much stronger *HPT* expression in the deletion lines lacking 21 nt sRNAs indicates that, indeed, they could be responsible for partial suppression, but against expectation, the plants remain hygromycin-resistant. In fact, while the role of 24 nt sRNAs in paramutation is becoming more and more obvious, the stability of *HPT* expression over more than 20 years and numerous rounds of seed amplification, despite the presence of the repeat and 21 nt sRNAs, is becoming the bigger enigma.

Another open question why Δ and γ deletions differ in their effect on the upstream promoter. We suppose that the difference lies probably not in the deletions themselves but in the way they were obtained. The Δ alleles originate from a T-DNA insertion mutagenesis, and although the T-DNA was not integrated, changes in chromatin states or modified interactions with other genomic regions are not uncommon outside of the T-DNA sites [67, 68]. We are not aware of similar effects after CRISPR mutagenesis. In addition, the Δ mutants were found after a rigorous hygromycin selection among thousands of lines, while CRISPR mutagenesis did not need to include a selection step.

These plants have been propagated in different locations, over decades, and under a variety of environmental conditions. Among thousands of scored seedlings of the R and S lines, we never observed a spontaneous switch of the epialleles from silent to active or from active to silent state. The latter might have gone undetected but certainly remains below detection. However, the temperature dependence of paramutation indicates that, after combination of the epialleles in the same plant, external factors can become highly relevant, with striking similarity to earlier observations with the *r1* paramutation system in maize [24]. However, involvement of small RNAs was not described in this maize system, and the expression of the *R* gene itself is regulated by light and temperature [24, 69], which is not the case for the *HPT* gene. A more detailed analysis of the temperature effects on paramutation should include sRNA libraries generated from different tissues and developmental stages but must consider different growth speed and the resulting asynchronous developmental stages. Temperature could also have a direct effect on the RNA by modifying secondary structures, probably less for the sRNAs but likely for the precursor and its processing by RDRs and Dicers. All we can say for now is that the prevention of paramutation at lower temperatures resembles the inhibition of RNAi-based viral defense in the cold [70], and the enhanced paramutation at higher temperatures supports an important role for the quantity of 24 nt sRNAs whose production is enhanced at higher temperature in both plants [71] and Paramecium, resulting in heterochromatin formation [72].

In summary, the data presented here for the *HPT* epialleles suggest that the occurrence and degree of paramutation are determined by multiple factors. The dependence on functional RdDM, the involvement of repeats, and the influence of environmental conditions overlap with features described for other paramutation phenomena. This study revealed an additional role for the copy number of the paramutable allele in the polyploid configuration and determination of paramutation through a delicate balance between the nature and amount of full length transcripts and small RNAs. The example of repeated switches between epigenetic states at the same locus, as reflected in the pedigree of the lines, provides evidence that spontaneous changes, targeted triggers, as well as genetic mutation events contribute to heritable stable yet reversible states and thereby to epigenetic diversity. With growing evidence for the existence of epialleles [73], it is tempting to speculate that paramutation between them often goes unnoticed and that paramutation may play an underestimated role in population genetics and evolution.

## MATERIAL AND METHODS

### Plant material

*Arabidopsis thaliana* ecotype Zürich (Zh) without (W) or with the transgenic epialleles (R and S) in diploid and tetraploid background were previously described [17]. Derivative lines (Δ) with genetic alterations leading to activation of the S epiallele were identified after failed T-DNA insertion [20]. RdDM mutants in tetraploid background and targeted deletion of the duplicated region were generated by CRISPR mutagenesis (see below). To generate tetraploid derivatives, 14 d-old diploid seedlings grown on plates were submerged in 0.1% colchicine (w/v) for 2 h, washed extensively in tap water and transferred to soil. Seeds from individual plants were harvested separately and sieved through a 350 µm mesh. Larger seeds retained in the sieve were more likely to be polyploid and were grown on soil to maturity. Ploidy of the progeny was evaluated by flow cytometry, by chopping approx. 100 mg of 14 d-old seedlings with a razor blade in 2 ml Galbraith buffer [74]. Nuclei were filtered through a 30 µm nylon mesh (CellTrics, Sysmex Partec, Görlitz, Germany) and pelleted by 10 min centrifugation at 4000 g and 4°C. The pellet was washed in 1 ml Galbraith buffer and centrifuged again. The pellet was resuspended in 500 µl Galbraith buffer with 5 µl of a solution containing 0.1 mg/ml 4’,6-diamidino-2-phenylindole (DAPI). After 10 min incubation on ice in the dark, samples were profiled for DAPI signal per nucleus on a BD LSRFortessa’ (BD Biosciences, Franklin Lakes, New Jersey, US) and compared to that of diploid and tetraploid control samples. An overview of all plant lines in this study is provided in Fig. 1.

### Growth conditions

Plants were regularly grown under long day (LD) conditions (16 h light 19°C, 8 h dark 16°C) and 60% relative humidity. For the temperature experiments with F1 hybrids, plants were grown on soil at standard conditions until development of the first inflorescences (approx. 21 days after germination) and then split into three cohorts: one remained in the standard conditions (19°C), the others were moved to chambers with either 10°C or 24°C under the same LD until seed set. Seeds were harvested, dried, and stratified at 4°C for 48 h before use.

### Crossings

All crosses were made after manual emasculation of immature flower buds and pollination by transfer of mature pollen onto the stigma of the prepared recipient. Contaminations were avoided by using sterilized tools and optical control under a binocular. All crosses were made in reciprocal orientation to exclude potential parent-of-origin effects.

### Hygromycin resistance assay

Seeds were aliquoted in 2 ml tubes and surface-sterilized for 10 min in a closed box with chlorine gas, generated by mixing 50 ml 14% sodium hypochlorite and 10 ml 37% hydrochloric acid. Seeds were sown on plates containing germination medium (GM, https://www.oeaw.ac.at/gmi/research/research-groups/ortrun-mittelsten-scheid/resources/) with or without 20 µg/ml hygromycin B (Calbiochem) and grown for 14 d under long day (LD) conditions (16 h light, 8 h dark 21°C). Plates were incubated horizontally or vertically for root growth assays. Resistance was evaluated by counting green plants with extended leaves and well-developed roots. Resistance ratios were calculated with reference to the number of all germinated seeds. Root length of plants grown on vertical plates was determined with Fiji (Version 1.51e) [75] and the corresponding SmartRoot plugin (Version 4.21) [76] relative to a ruler and size of the plate. Statistical analysis via Chi-square test was performed with the program R (Version 4.0.2; https://www.r-project.org/) or the Welch two sample T-test function for of heteroscedastic data in R. Raw data for these and all other assays are available in S2 Table.

### DNA and RNA extraction

For genotyping, DNA was extracted from one young leaf in a 2 ml tube containing approx. 10 glass beads and 500 µl extraction buffer (200 mM Tris pH 7.5, 250 mM NaCl, 25 mM EDTA) and ground in an MM400 homogenizer (Retsch, Düsseldorf, Germany) for 1 min at 30 Hz. Samples were centrifuged 1 min at 16000 g and 450 µl supernatant was transferred into a 1.5 ml tube containing 450 µl isopropanol and 50 µl sodium acetate pH 5.2. After >1 h at - 15 20°C, precipitates were collected by 20 min centrifugation at 16000 g, washed once in 75% EtOH and air-dried. Pellets were dissolved in 70 µl TE-buffer and stored at 4°C until usage.

For PCR analysis, DNA was prepared from 0.1 – 1 g plant material ground in liquid nitrogen and suspended in 0.3 – 3 ml CTAB elution buffer (A4150; PanReac AppliChem, Darmstadt, Germany, supplied with 1% polyvinyl pyrrolidone (PVP) and 1% β-mercaptoethanol (β-ME)). After adding RNase A (EN0531, Thermo Fisher Scientific, Waltham, Massachusetts, US), samples were shaken 2 h at 65°C followed by phenol-chloroform extraction as described [77]. DNA was precipitated as above and dissolved in an appropriate amount of TE buffer. Concentrations were determined on a Nanodrop ND1000 (Thermo Fisher Scientific, Waltham, Massachusetts, US). DNA samples were stored at -20°C.

RNA was extracted from either 100 mg 14 d-old seedlings or 30 mg 35 d-old flower buds with TRI-Reagent (T9424, Sigma Aldrich, St. Louis, Missouri, US) according to the manufacturer’s protocol. Total RNA was eluted in 50 µl RNase-free ddH2O and the concentration determined as for DNA. RNA integrity was analyzed by running aliquots on a 1.5% agarose-TAE gel, before treating 2.5 μg with DNaseI (EN0525, Thermo Fisher Scientific, Waltham, Massachusetts, US) according to the manual. DNAseI was heat-inactivated at 65°C for 10 min in the presence of 10 mM EDTA. DNA-free total RNA (RNA+) was precipitated by adding 1/10 volume sodium acetate pH 5.2 and 3 volumes 100% cold EtOH for > 2 h at -20°C. RNA was pelleted and washed once in 700 µl cold 75% EtOH. Air-dried pellets were dissolved in 40 µl RNase-free ddH2O and stored at -70°C until use.

### Quantitative RT-PCR

cDNA was synthesized from total RNA with RevertAid H Minus Reverse Transcriptase (EP0451; Thermo Fisher Scientific, Waltham, Massachusetts, US), with Random Hexamer primers (S0142; Thermo Fisher Scientific, Waltham, Massachusetts, US) according to the manufacturer’s recommendation, but extending cDNA synthesis at 42°C for 90 min in the presence of 20 U RiboLock RNase Inhibitor (EO0381; Thermo Scientific). Prior to quantitative real-time PCR (qRT-PCR), absence of genomic DNA contaminations was controlled by 40 cycles PCR without reverse transcriptase (RT) at 60°C annealing and 1 min elongation, or with RT and primers spanning the intron of the UBC28 gene (At1g64230), in 30 cycles with 20 sec elongation. qRT-PCR was performed with FastStart Essential DNA Green Master (Roche; Rotkreuz, Switzerland) with approx. 5 ng of cDNA on a LightCycler96 system (Roche) according to the manufacturer’s instructions. Amplification was performed in a two-step protocol with pre-incubation at 95°C for 10 min and 45 cycles alternating between 95°C 10 sec/60°C 30 sec. A final melting cycle at 97°C was done preceding melting curve analysis. Data analysis was performed according to the ΔΔCt method [78] relative to the housekeeping genes *SAND* (At2g28390, Fig 5C) or *EIF4A1* (At3g13920, Fig 6B). Fold change values relative to the control RR or RRRR, respectively, are shown. Statistical analysis was performed with the R-package “agricolae” (Version 1.3-3; https://cran.r-project.org/web/packages/agricolae/index.html) applying a one-way ANOVA followed by a post-hoc Tukey HSD test (α = 0.05). The following primers were used: fw 5’ GGGTAAATAGCTGCGCCGATGGTT and rv 5’ CACGGCGGGAGATGCAATAGGTC (expression of *HPT*); fw 5’ AACTCTATGCAGCATTTGATCCACT and rv 5’ TGATTGCATATCTTTATCGCCATC (qRT-PCR normalization to *SAND*/At2g28390); fw 5’ ATCCAAGTTGGTGTGTTCTCC and rv 5’ GAGTGTCTCGAGCTTCCACTC (qRT-PCR normalization to *EIF4a*/At3g13920). fw 5’ TCCAGAAGGATCCTCCAACTTCCTGCAGT and rv 5’ ATGGTTACGAGAAAGACACCGCCTGAATA (spanning a small intron in *UBC28*/At1g64230 to exclude contamination of cDNA with genomic DNA).

### Northern blotting

Gels with 5 µg total RNA per slot were blotted onto Hybond NX membranes (Amersham, UK) by capillary blotting with 20 x SSC overnight. Membranes were washed once in 2 x SSC, air- dried and crosslinked in a UV Stratalinker 2400 (Stratagene, La Jolla, California, US) in Autocrosslink mode. To visualize size markers and to confirm equal loading, crosslinked membranes (DNA and mRNA blots) were stained for 5 min at RT with methylene blue staining solution (0.04% methylene blue [w/v] in 0.5 M sodium acetate pH 5.2) and washed in ddH2O. Methylene blue stain was removed by incubation at RT for 20 min in 2 x SSC with 1% SDS on a shaker and then air-dried.

Probes for nucleic acid blots were generated by PCR amplification of the corresponding region of template DNA or made from oligonucleotides.. Amplicons were gel-purified (D4002; Zymo, Irvine, California, US) and diluted to 25 ng/µl. Twenty-five ng were labelled with 50 µCi [α- 32P]dCTP (SRP-305; Hartmann Analytic; Braunschweig; Germany) with the Amersham Rediprime II Random Prime Labeling System (RPN1633; GE Healthcare, Chalfont St Giles, UK) and purified on Illustra G-50 ProbeQuant micro columns (GE Healthcare, Chalfont St Giles, UK). Prior to hybridization, the longer, double-stranded probes were denatured for 5 min at 99°C. These probes were used for the detection of HPT transcripts on northern blots and the hairpin-derived sRNAs. For the detection of endogenous sRNAs such as siRNA1003 (siR1003_probe: 5’ ATGCCTATGTTGGCCTCACGGTCT) and miR160 (miR160_probe: 5’ tggcatacagggagccaggca), twenty-five mM oligonucleotide were end-labelled with T4 Polynucleotide Kinase (PNK; EK0031; Thermo Scientific) according to the manual with 25 µCi [γ-32P]ATP (SRP-501; Hartmann Analytic; Braunschweig; Germany) for 45 min at 37°C. Radioactive labelled oligonucleotide probes were purified with Illustra MicroSpin G-25 micro columns (GE Healthcare, Chalfont St Giles, UK).

Blots were hybridized as described [79] in a hybridization oven overnight at 42°C for oligo- probes or 60°C for longer probes. Membranes were washed twice for 20 min each in 2 x SSC + 2% SDS at 50°C for oligo-probes or 65°C for longer probes, followed by 15 min in 2 x SSC +0.2% SDS at the respective temperatures. Membranes were wrapped in Saran wrap and exposed to phosphoscreens (GE Healthcare, Chalfont St Giles, UK) for up to 3 days. Screens were scanned on a phosphoimager (Typhoon FLA 9500, GE Healthcare, Chalfont St Giles, UK).

### Preparation and sequencing of sRNAs

Sequencing libraries were prepared from either 500 ng total RNA or 3 µl TraPR-enriched [80] sRNA with the NEBNext Multiplex Small RNA Library Prep Set for Illumina (E7300; NEB, Ipswich, Massachusetts) according to the manufacturer’s protocol. Libraries were amplified in 14 cycles and cleaned up with AMPure XP beads (Beckman Coulter, Brea, California, US) prior to size selection on 6% polyacrylamide gels. cDNA libraries from approx. 140 bp to 150 bp (miRNAs/siRNAs including adapters) were gel-eluted as described in the manufacturer’s protocol, EtOH-precipitated at -20°C overnight and eluted in 12 µl TE buffer. Size range of the libraries was confirmed on a Fragment Analyzer (Agilent formerly Advanced Analytical, Santa Clara, California, US) with the HS NGS Fragment Kit (1-6000 bp, DNF-474, Agilent, formerly Advanced Analytical, Santa Clara, California, US). sRNAs were sequenced on a HiSeq 2500 system in single end 50 bp mode (Illumina, San Diego, Santa Clara, California, US) by the VBCF NGS facility. Statistics for the libraries is summarized in S1 Table.

### Plasmid construction and plant transformation

Cloning for subsequent Sanger sequencing and subcloning steps was made with the CloneJET PCR cloning kit (K1231, Thermo Fisher Scientific, Waltham, Massachusetts, US). For plant transformation with the recloned epiallele, we used a pSUN vector containing an oleosin promoter upstream of GFP [81]. For Cas9-expressing vectors, the ubiquitin4-2 promoter (PcUbi4-2) of *Petroselinum crispum* in the binary vector pDE-Cas9 [33] was replaced with the egg cell-specific promoter EC1.2p [82] and the previously described seed-specific GFP marker was inserted to generate the pDEECO vector backbone. Two specific sgRNAs each were expressed from an Arabidopsis U6-26 promoter and processed via the tRNA-based multiplex system [83]. For ectopic silencing constructs, the PPT cassette of pDE-Cas9 containing a minimal 35S promoter sequence was removed, and the SpCas9 coding sequence was replaced by the PDK/catalase intron sequence with flanking multiple cloning sites from the pHELLSGATE12 vector [84], resulting in the pDEPPi backbone. Sequences matching different regions of the epiallele were inserted in sense and antisense direction to result in hairpin-forming transcripts. All vectors were transformed into E. coli strains DH5α. Plasmids were prepared with Zyppy Plasmid Miniprep Kit (D4020, Zymo, Irvine, California, US) and eluted in 50 µl ddH2O. Plasmids were controlled by Sanger sequencing before being transformed into electrocompetent *Agrobacterium tumefaciens* strain GV3101 pMP90. Arabidopsis plants were transformed with the floral dip method [85] and transformants selected among seeds of the T0 plants based on the GFP signal in seeds as described [81].

### CRISPR mutagenesis

sgRNAs were designed with Chopchop [86–88] and ranked based on their efficiency score and suitable target sites. Prior to cloning the vectors for deletion mutagenesis, we tested several sgRNA candidates for efficient cleavage at the target site in an *in vitro* assay, incubating reconstituted sgRNA-Cas9 ribonucleoprotein with amplicons containing the target site [89]. For genotyping potential mutant candidates, the *in vitro* assay was modified by replacing the wild type amplicon with that of pooled genomic DNA from candidate plants [89]. PCR products were analyzed by electrophoresis, and the analysis repeated for individual plants in positive pools. Amplicons around the mutated site were further analyzed by Sanger sequencing to determine the type of mutation.

### sRNA mapping and profiling

From the reads of the sRNAs libraries, adapters were trimmed and reads were size-selected to the length of 18 nt–26 nt by using cutadapt (Version 1.9.1; https://doi.org/10.14806/ej.17.1.200), with options -a AGATCGGA-g CGACGATC -m 18 -M 26 -discard-untrimmed. Read quality and size distribution was controlled using FastQC (Version 0.11.5; http://www.bioinformatics.babraham.ac.uk/projects/fastqc/). Reads were aligned to the TAIR10 genome [90] including an extra contig containing the sequence of the transgenic epiallele plus upstream and downstream flanking regions. Alignment was done using Bowtie2 [91] with the options -k 500 -no-unal.

Annotation was done by comparison of genomic positions of small RNA sequences using the Araport11 and repeat-masker databases. For small RNAs with multiple genomic locations, an annotation was attributed given a priority order.

For small RNA profiles, reads aligned to the transgenic epiallele and flanking regions were retrieved and used to calculate the normalized read counts (reads per millions mapped reads) for each nucleotide and to produce a graphical representation using R.

A complete list of procedures and scripts used for bioinformatic analysis and graphical representation are available on GitHub (https://github.com/AlexSaraz1/paramut_bot).

sRNA sequencing data are deposited in the Gene Expression Omnibus (GEO) database (https://www.ncbi.nlm.nih.gov/geo/) under the accession number GSE162241 and raw data in the Sequencing Read Archive (SRA, https://www.ncbi.nlm.nih.gov/sra) under the number SRP294483. An overview over the sRNA libraries and their GEO accession numbers have been provided in S1 Table.

## ACKNOWLEDGEMENTS

We are grateful for helpful discussions within our group and colleagues on campus, especially Arturo Marí-Ordóñez, Peter Schlögelhofer, and Michael Nodine. We thank the lab of Olivier Voinnet for suggestions and discussions, and in particular Thomas Grentzinger for supplying the TraPR columns used for sRNA purification ahead of publication and Alexis Sarazin who provided an in-house sRNA-sequencing data processing pipeline. We are also grateful to Helene Fasching and Emi Miyakoda for help with resistance assays and genotyping. We thank the campus facilities, especially Next Generation Sequencing and Plant Facilities from the VBCF and MolBiol Service and BioOptics from the IMP/IMBA/GMI facilities for their excellent support. We thank Arturo Marí-Ordóñez, Liam Dolan, Vincent Castric, J. Matthew Watson, and three anonymous reviewers for valuable comments on the manuscript.

## SUPPORTING FIGURE LEGENDS

**S1 Figure.**
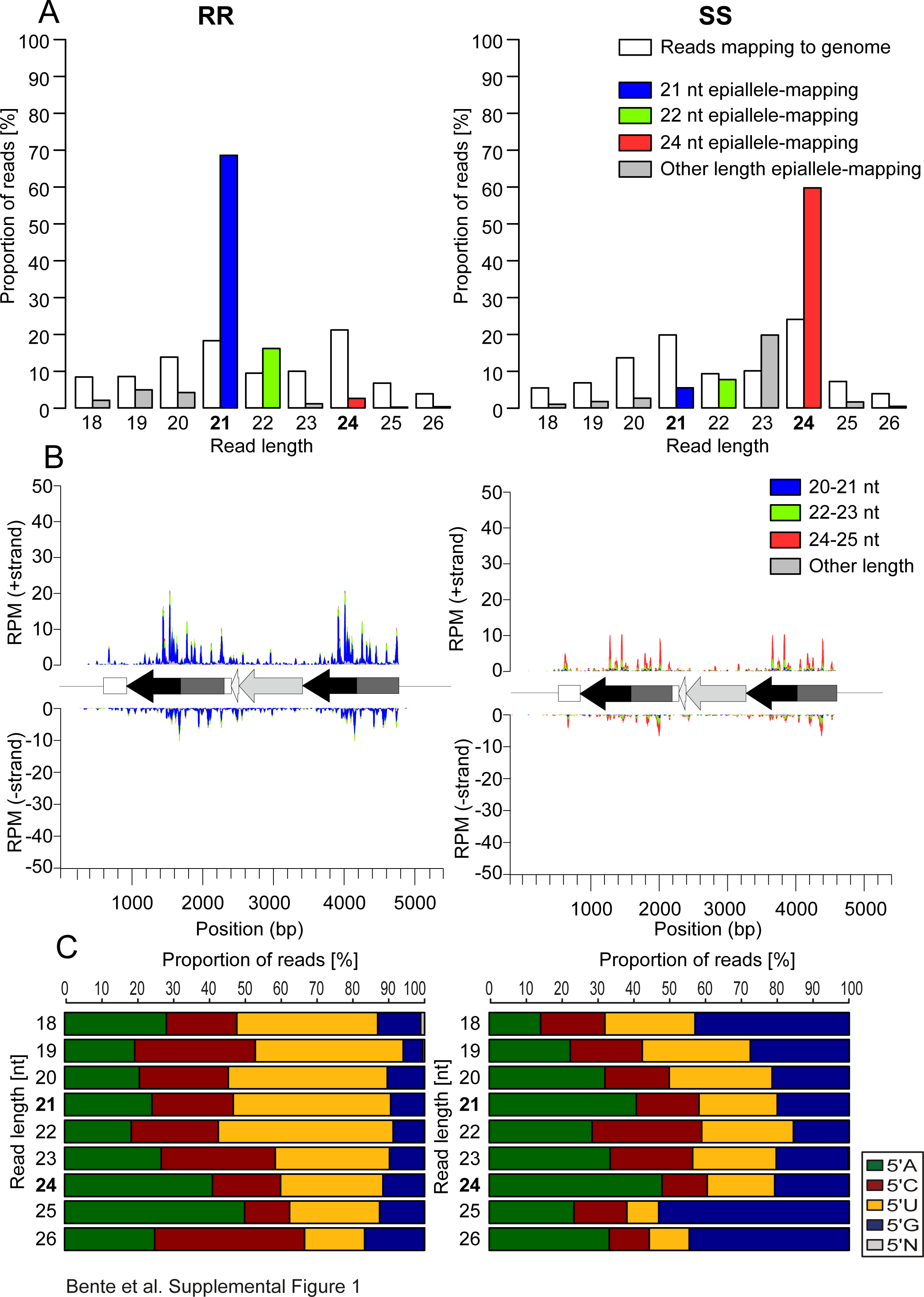
Epialleles associated with distinct classes of sRNAs in diploids. (A) Proportion of sRNA length from all mapped reads in 14 day-old seedlings with diploid active (RR) or silenced (SS) epialleles. (B) Coverage plots of 18 - 26 nt sRNA along the RR or SS epialleles. (C) Proportion of 5’ prime nucleotides of epiallele-specific sRNAs in RR (left) and SS (right) plotted by size.

**S2 Figure.**
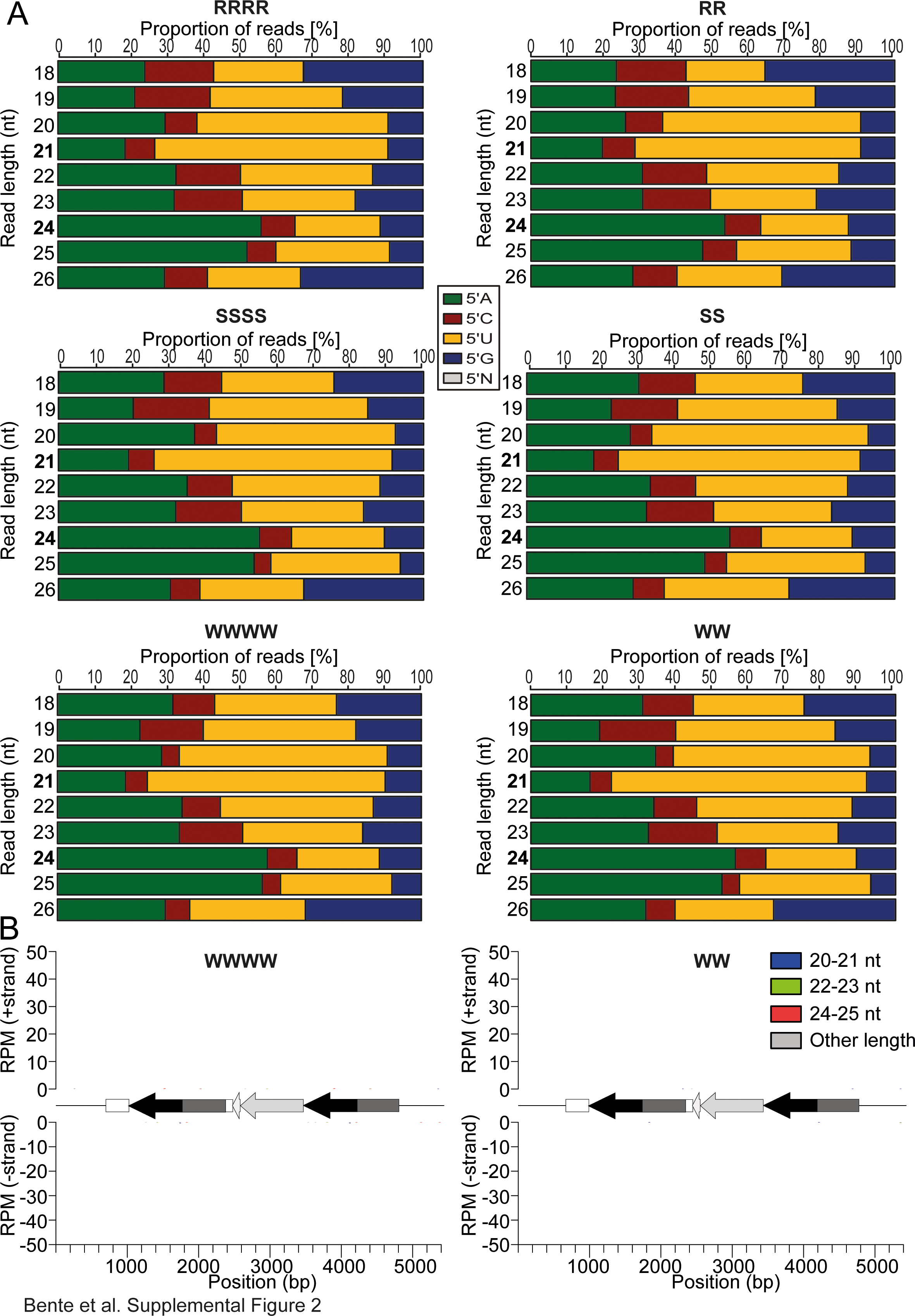
Equal sRNA distribution genome-wide and no sRNAs mapping to the epiallele sequence in wild type. (A) Proportions of 5’ nucleotides in different size classes in sRNA libraries from tetraploids (left) and diploids (right) in lines with active epialleles (top), silent epialleles (middle), and wildtype lines (bottom). (B) RNA libraries from tetraploid (WWWW, left) or diploid (WW, right) plants were attempted to map to the full-length sequence of the epiallele.

**S3 Figure.**
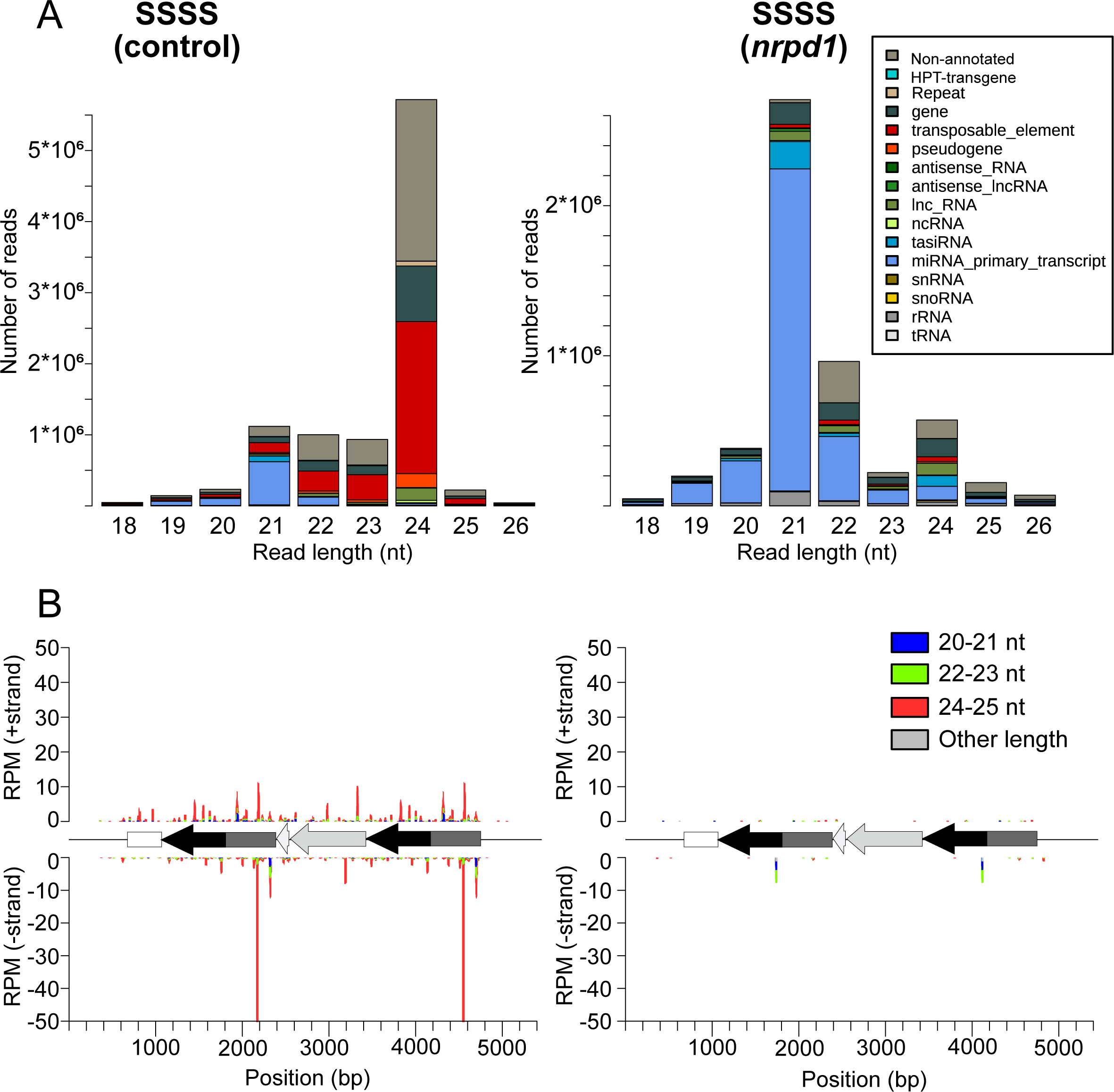
Effects of the Cas9-generated PolIV mutant on sRNA profiles. (A) Strong reduction of 24 nt sRNAs in a homozygous loss-of-function mutation in the *NRPD1* gene (encoding the largest subunit of PolIV) generated by CRISPR in the background of the tetraploid silent epiallele; including (B) those mapping to the epiallele.

**S4 Figure.**
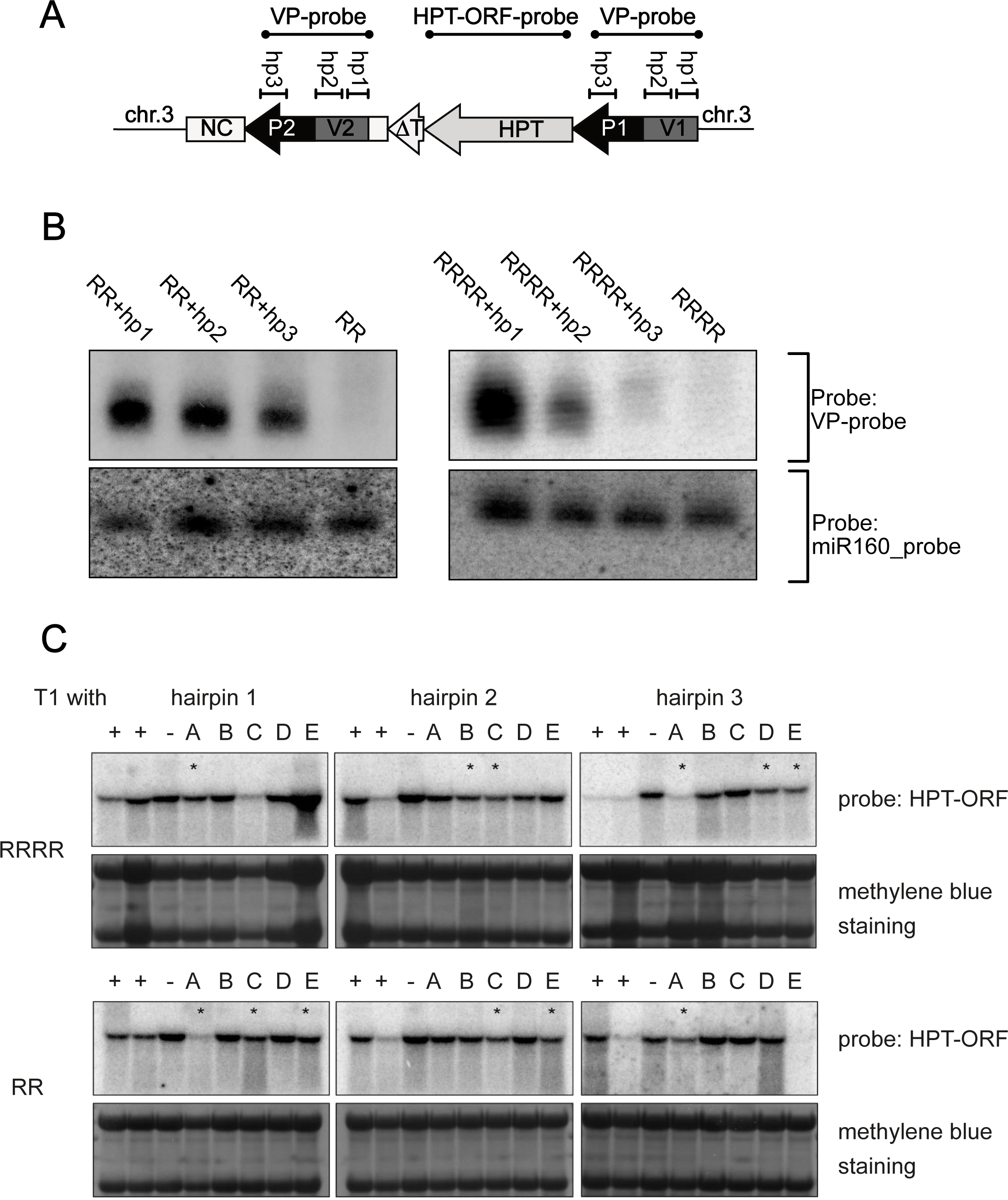
sRNAs produced by the hairpin constructs and lasting effects on HPT expression. (A) Location of the hairpin sequences and region covered by the VP- and HPT- ORF probe for detection in B and C. (B) Northern blots with sRNA from flower buds of 35 d-old diploid and tetraploid T1 plants selected to contain the hairpin constructs. VP probe (labelled by random priming, top) and antisense miR160 (end-labelled oligonucleotide) as loading control (bottom). (C) Northern blot analysis of HPT transcript from tetraploid (top) or diploid (bottom) plants with R epialleles never containing the hairpin (-), individual T2 plants containing the hairpin (+) or not containing it any more due to segregation (A-E). Five µg of total flower bud RNA from 35 d-old plants was hybridized with the HPT-ORF or stained with methylene blue as loading control. Asterisks mark samples with reduced HPT transcript despite the absence of the hairpin.

**S5 Figure.**
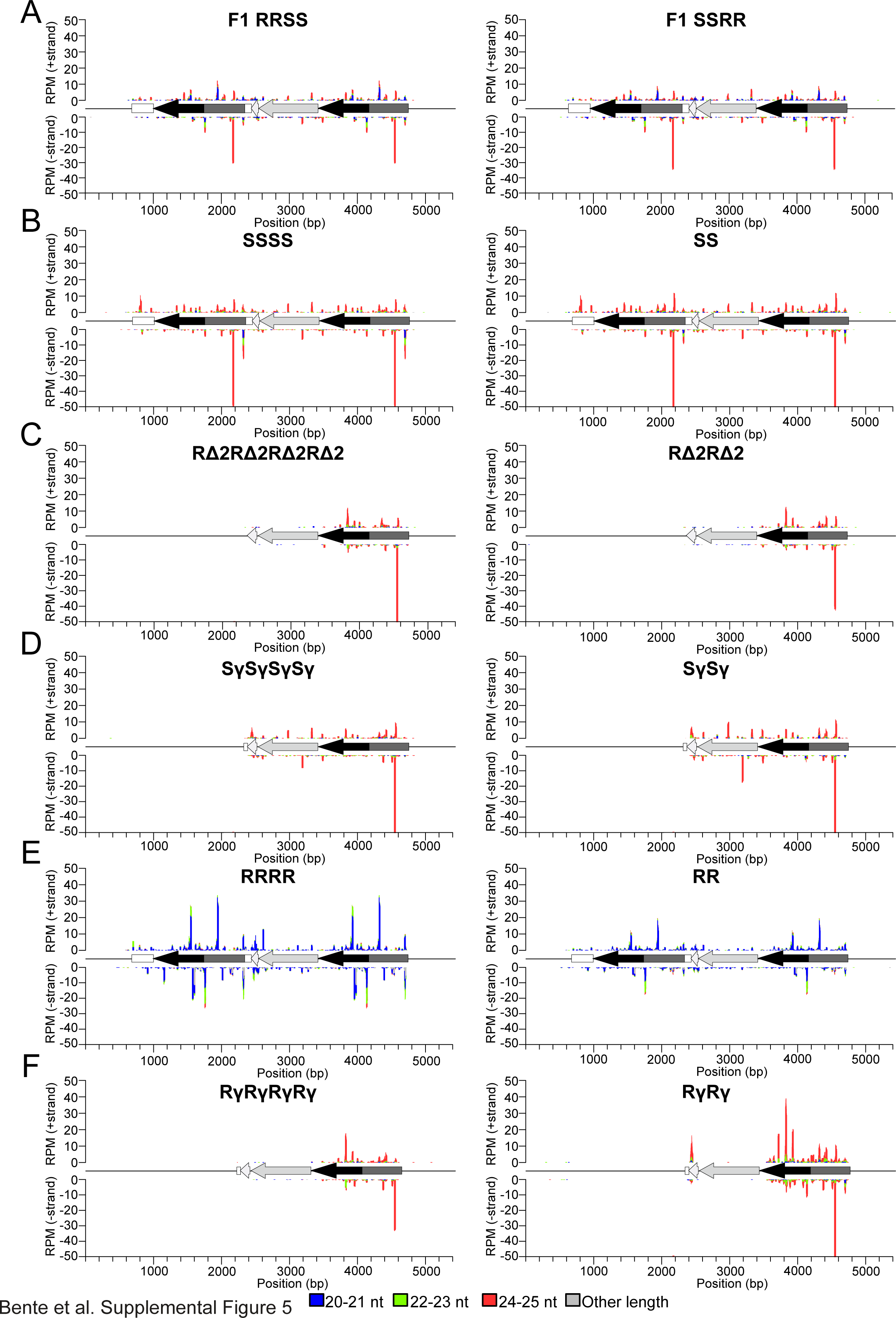
sRNA profiles along the epialleles in F1 hybrids and in deletion mutants. (A) Profile from flower bud material in a pair of reciprocal paramutation test hybrids. (B – F) Profiles of tetraploid (left) and diploid (right) lines with the indicated alleles; (B) silent full length allele; (C) active allele missing downstream repeat after random mutagenesis and hygromycin screen; (D) CRISPR-generated deletion of the downstream repeat in the background of the silent epiallele; (E) active full-length allele; (F) CRISPR-generated deletion of the downstream repeat in the background of the active epiallele. All values are reads per million mapped reads (RPM), and the y-axes are all set to the same maximum of ± 50 RPM.

**S6 Figure.**
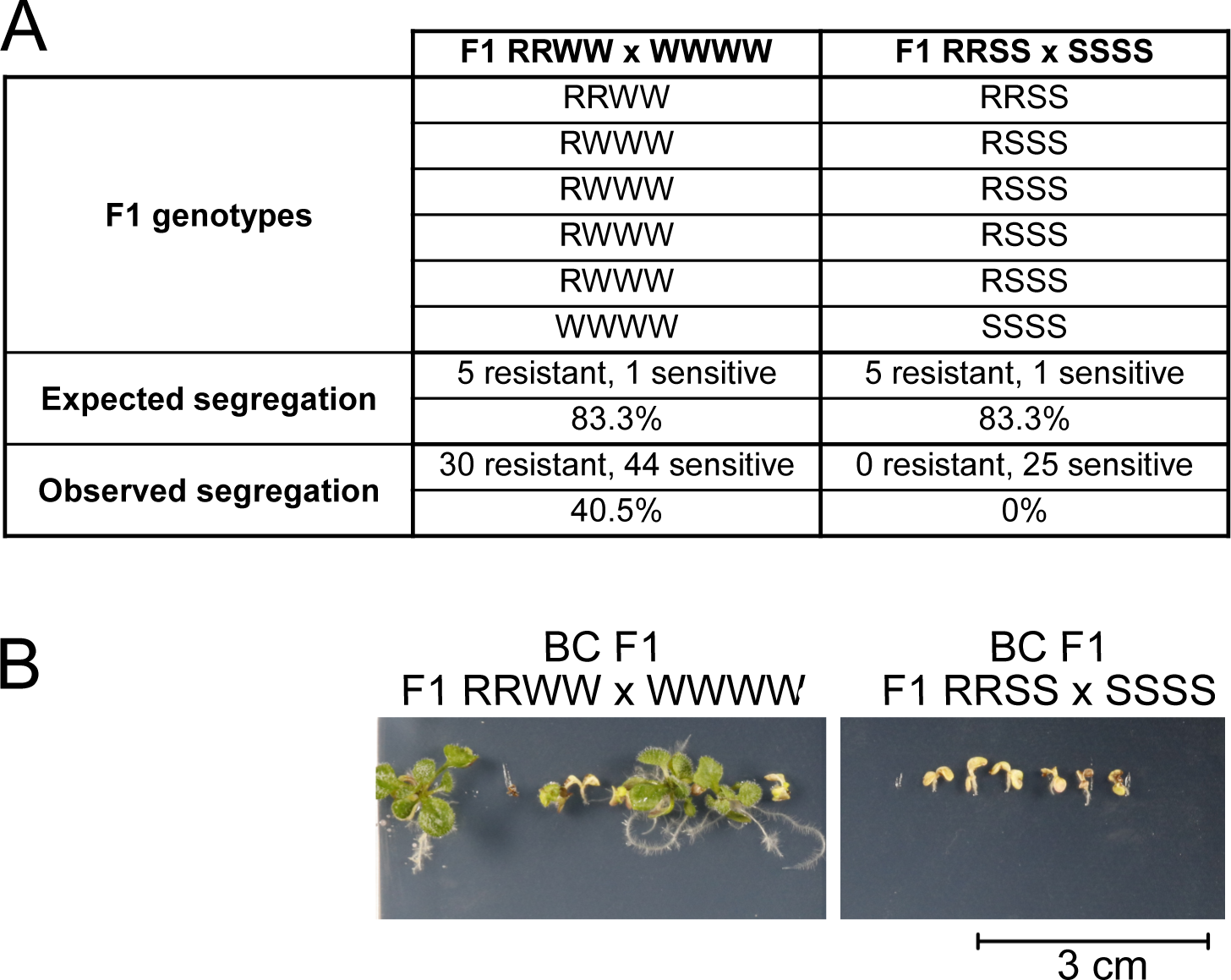
Supporting data for the dosage effect. (A) Tetraploid hybrids with two alleles each of R and W, or R and S, were backcrossed to homozygous W or S, respectively. In both cases, two thirds of the F1 progeny will contain one R epiallele, and 5 out of 6 plants can be expected to be resistant. The lack of any resistant plants in the crosses involving S indicates a strong silencing effect in case of three S in combination with one S. (B) Representative picture of three-week-old plants from the indicated genotypes grown on hygromycin selection medium. Scale bar = 3 cm.

S1 Table. Numerical data for the sRNA libraries.

S2 Table. Numerical data for hygromycin resistance assays, expression analyses, and root growth measurements. Each tab provides the numerical data and basic statistics for one figure in this study.

